# Structural mechanism underpinning *Thermus oshimai* Pif1-mediated G-quadruplex unfolding

**DOI:** 10.1101/2021.08.24.457465

**Authors:** Yang-Xue Dai, Hai-Lei Guo, Na-Nv Liu, Wei-Fei Chen, Stephane Rety, Xu-Guang Xi

**Affiliations:** College of Life Sciences, Northwest A&F University, Yangling, Shaanxi 712100, China; Université de Lyon, ENS de Lyon, Université Claude Bernard, CNRS UMR 5239, INSERM U1210, LBMC, 46 allée d’Italie, Site Jacques Monod, 69007, Lyon, France; Laboratoire de Biologie et de Pharmacologie Appliquée (LBPA), UMR 8113 CNRS, Institut d’Alembert, Ecole Normale Supérieure Paris-Saclay, Université Paris-Saclay, 4, Avenue des Sciences, 91190 Gif sur Yvette, France

## Abstract

G-quadruplexes (G4s) are unusual DNA structures and can stall DNA replication, causing genomic instability for the cell. Although the solved crystal structure of the DHX36 helicase demonstrated that G4 was specifically targeted by a DHX36-specific motif (DSM), lack of complete structural details for general G4-resolving helicases without specific target motifs remains a barrier to the complete understanding of the molecular basis underlying the recognition and unfolding of G4s. Herein, we present the first X-ray crystal structure of the *Thermus oshimai* Pif1 (*To*Pif) complexed with a G4, thereby mimicking the physiological G4 formed during DNA replication. Strictly different from the previous determined G4-helicase structure of DHX36, our structure revealed that *To*Pif1 recognizes the entire native G4 via a cluster of amino acids at domains 1B/2B constituting a G4-Recognizing Surface (GRS). The overall topology of the G4 structure solved in this work maintains its three-layered propeller-type G4 topology, with no significant reorganization of G-tetrads upon protein binding. The three G-tetrads in G4 were differentially recognized by GRS residues mainly through electrostatic, ionic interactions and hydrogen bonds formed between the GRS residues and the ribose-phosphate backbone. Our structure explains how helicases from distinct superfamilies adopt different strategies for recognizing and unfolding G4s.

## INTRODUCTION

G-quadruplexes (G4s) are higher-order DNA/RNA structures formed from nucleic acid sequences that contain four stretches of three or more guanines interspaced by at least one random nucleotide (1,2). The G-tetrad module, composed of four guanines, forms a planar structure stabilized by non-canonical Hoogsteen hydrogen bonds in the presence of monovalent cations, such as sodium or potassium (2,3). Genome-wide bioinformatics analyses have revealed the prevalence of G4 motifs in key regulatory regions of the human genome, such as promoters, gene bodies and untranslated regions (4–8). In accordance with this prediction, a tremendous increase in cell experimental data has demonstrated that G4 structures do indeed exist under physiological conditions and are involved in a variety of biological processes including telomere maintenance, gene expression, epigenetic regulation and DNA replication (9,10). The high thermodynamic stability of G4s in cells can erect significant barriers to replication fork progression. Therefore, failing to resolve the formed G4s during DNA transactions can cause replication to stall and thus may trigger the rampant genomic instability characteristic of certain cancer cells (11–14). Cells have evolved the capability of resolving stable G4 with a group of helicases that play important roles in the maintenance of genomic stability in many organisms (15). The G4-resolving helicases are found in most helicase superfamilies (SF), including SF1 and 2 (SF1: e.g. Pif1; SF2: e.g. DHX36 and the RecQ family of helicases (BLM, WRN, RecQ)).

The DHX36 protein, one of most extensively studied G4-unfolding helicases, is characterized by DHX36-specific motif (DSM) and specifically binds parallel G4s. Surprisingly, the X-ray crystal structure of DHX36 (16) in complex with a Myc-promoter-derived G4 demonstrates a rearrangement of the 5′-G-tetrad with a one-base translocation upon DSM binding (16). Because a NMR study shows that G4 keeps its integral state upon the binding of an isolated DSM (17), it was therefore suggested that DSM, in coordination with the helicase core, unfolds the G4 one base at a time in an ATP-hydrolysis-independent manner. Recently, a crystal structure (18) solved from a helicase core of a RecQ family helicase (*Cs*RecQ) complexed with oligonucleotides harboring a human telomere sequence revealed that a guanine base within the putative G-tetrad is located at the 5′-gate of the electropositive channel where the G-base is flipped and sequestrated (18). This structure is interpreted to represent “a product complex of *Cs*RecQ-unwound G4 rather than a folded quadruplex (18)” although the complexed single-stranded DNA (ssDNA) was not prefolded to form a stable G4 prior to the preparation of the G4-*Cs*RecQ complex. Crystal structures of Pif1 helicases from Human, *Saccharomyces cerevisiae, Bacteroides sp* and *Thermus oshimai* have been recently solved (19–22). Pif1 is a helicase conserved from bacteria to humans, and has been shown to unfold G4 and to be critical for genome stability in yeast (23–25). A recent study has demonstrated that PIF1 helicase promotes break-induced replication in mammalian cells (26).

Elucidating the molecular mechanism by which G4-resolving helicases recognize and unfold G4 molecules largely depend on the knowledge of crystal structures. Access to structures representing a “substrate” complex, intermediates and a “product” complex provides very valuable mechanistic information. Although the helicase-G4 DNA complex structures of DHX36 (16) and *Cs*RecQ (18) can be considered as representations of an intermediate substrate complex (16) and a final product complex of helicases (18), respectively, the remaining challenge is to obtain the structure in which the helicases interact with an intact G4 (initial substrate complex) or to capture the structure of the very first steps leading to transition state.

In this study, we report the X-ray crystal structures solved from the Pif1-family helicase from *T. oshimai* (*To*Pif1) bound to a prefolded G4 bearing 6-8 nt on both sides of the G-tetrads, thereby mimicking the physiological G4 state formed during DNA replication. This is the first structural snapshot of a helicase complexed with an integral G4. In combination with bulk and single molecular fluorescent resonance energy transfer assays (smFRET), our studies provide new mechanistic insight into how a Pif1-family helicase recognizes, binds and unfolds G4s.

## MATERIALS AND METHODS

### Reagents and buffers

All chemicals were reagent grade and all buffers were prepared in high-quality deionized water from a Milli-Q ultrapure water purification system (Millipore) having resistivity of >18.2 MΩ.cm and were filtered again on a 0.20 μm filter before use. All DNA unwinding and binding assays were performed in buffer A (20 mM Tris-HCl (pH 7.5 at 37 °C), 50 mM NaCl, 2 mM MgCl_2_, 2 mM DTT) which was optimized previously by our group.

### DNA substrate preparation

All the DNA substrates used in this study were chemically synthesized and HPLC-purified by Sangon Biotech (Shanghai) and are listed in Supplementary Table S1. The oligonucleotides used in binding and unwinding assays were prepared at a 2 μM working concentration. The duplex or G4 DNA used in the stopped-flow and smFRET assays were heated to 95 °C for 5 min in stocking buffer (20 mM Tris-HCl, pH 7.5, 100 mM NaCl) and annealed by slow cooling to room temperature. The DNA substrates in crystallization were dissolved in 20 mM Tris-HCl (pH 7.5) with 100 mM NaCl, heated to 95 °C, and allowed to cool slowly to room temperature in a water bath. After purification on a Mono Q column, the formation of G4 structures was checked using circular dichroism (CD) spectropolarimetry.

### Protein expression and purification

*To*Pif1 (residues 64-507) and its mutants (R355A, R135A, R150A, R419A, R392A, Q327A, K329A, E397A, E397L, E397H and E397D) were all cloned into pET15b-SUMO and then transformed into the C2566H *E. coli* strain (New England Biolabs), respectively. When the culture reached early stationary phase (OD_600_ = 0.55 - 0.6) at 37 °C, 0.3 mM IPTG was added and the protein expression was induced at 18 °C over 16 h. Cells were harvested by centrifugation (4500 g, 4 °C, 15 min) and pellets were suspended in lysis buffer (20 mM Tris-HCl pH 7.5, 500 mM NaCl, 10 mM Imidazole and 5% glycerol (v/v)). Cells were broken with a French press and then further sonicated 2-3 times to shear DNA. After centrifugation at 12,000 rpm for 40 min, the supernatants were filtered through a 0.45-μm filter and loaded onto a Ni^2+^ charged IMAC column (GE Healthcare). After washing twice, the SUMO-ToPif1 was then eluted from the Ni^2+^ affinity column with elution buffer (20 mM Tris-HCl, pH 7.5, 500 mM NaCl, 300 mM Imidazole and 5% glycerol (v/v)) at 4 °C. The eluted protein was treated with SUMO protease (Invitrogen, Beijing). The eluted protein was treated with SUMO protease (Invitrogen, Beijing). Then the SUMO digested protein was further purified by a HiTrap Heparin column (GE Healthcare) to remove the SUMO-tag and other protein impurity. The eluted fraction containing ToPif1 was collected and concentrated. The final purified protein was dialyzed against the storage buffer (20 mM Tris-HCl, pH 7.5, 500 mM NaCl, 1 mM DTT) and concentrated to approximately 10 mg/mL for crystallization and was about 95% pure as determined by SDS-PAGE. Mutations and truncations were engineered by PCR overlapping-PCR protocol.

### Equilibrium binding assays

The isothermal binding curves were determined using a fluorescence polarization assay on an Infinite F200 plate reader (Tecan). FAM-labeled DNA substrates were used in this study. Varying amounts of protein were added to a 150 μL aliquot of buffer A containing 5 nM FAM-labeled DNA. Each sample was allowed to equilibrate in solution for 5 min at 37 °C, and then fluorescence polarization was measured. Less than 5% change was observed between the 5 and 10 min measurements, indicating that equilibrium was reached in 5 min. The equilibrium dissociation constants were determined by fitting the binding curves using Equation 1:

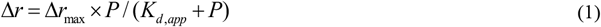

where Δ*r*_max_ is the maximal amplitude of the anisotropy (i.e., *r*_max_ – *r*_free,DNA_), *P* is the helicase concentration, and *K*_d,app_ is the midpoint of the curve corresponding to the apparent dissociation constant.

### Crystallization of *To*Pif1-nucleic acid complexes

For crystallization, purified *To*Pif1 was incubated with the G4^T6/T8^ at a molar ratio of 2:1 in the presence of ATP analog ADP·AlF_4_. The resulting *To*Pif1-G4^T6/T8^-ADP·AlF_4_ complex was purified using size-exclusion chromatography and then concentrated to approximately 10 mg/mL. Crystallization trials on *To*Pif1 and its complexes with DNA (Supplementary Table S1) and ADP·AlF_4_ were performed at 20 °C using the sitting-drop vapor diffusion method. Crystallization screening was carried out at 20 °C using commercial screening kits (Hampton Research, Molecular Dimensions and Rigaku Reagents), where the *To*Pif1-DNA complex was mixed at a 1:1 ratio with the reservoir solution. Crystals of *To*Pif1-G4^T6/T8^-ADP·AlF_4_ were obtained using 0.1 M Bis-Tris (pH = 6.5), 10% PEG20000. All these conditions were optimized with a grid search using 48-well Linbro plates at 20 °C where 1 μL of protein sample and 1 μL of precipitant were mixed together and equilibrated with 60 μL of precipitant.

### X-ray data collection, phasing and refinement

All X-ray diffraction data were collected on beamline BL19U1 (27) at the Shanghai Synchrotron Radiation Facility (China) using a Pilatus 6M detector (Dectris) and were processed using XDS (28). *To*Pif1-G4^T6/T8^-ADP·AlF_4_ structures were solved by molecular replacement, performed in the PHENIX software suite (29) with Phaser (30), using the *To*Pif1 apo structure (PDB: 6S3E (20)) as the template model (31). Manual reconstruction was done using the Coot (32) and further refinement was performed in PHENIX. Cell parameters and data collection statistics are reported in Supplementary Table S2.

### Small-angle X-ray scattering assay

SEC-SAXS experiment was carried out at beamline SWING (SOLEIL Synchrotron, Saint-Aubin, France) with SEC-HPLC coupled to SAXS. The sample of ToPif1-G4 complex was injected at a concentration of 10 mg/ml on a Superdex 200 5/150 Increase column (Cytiva) at a flow rate of 0.2 ml/min equilibrated in buffer 25 mM Hepes (pH=7.5), 150 mM KCl, 5%glycerol. Scattering data were collected at 20 °C usinga Pilatus 1M dectector (Dectris) and data reduction and processing of images were done with Foxtrot (33). Further analysis was done with ATSAS 2.8 suite (34). Experimental *R*_g_, *I*(0), and *D*_max_ values were calculated with PRIMUS and GNOM4 programs respectively. *Ab initio* envelopes for isolated complexes were determined using DAMMIF with the pair distance distribution function calculated with GNOM4. Full atomic models derived from crystal structure was modelled with MODELLER and adjusted to SAXS data with DADIMODO. In this modelling procedure, only the missing parts from X-ray structure were kept flexible. Profiles of atomic models were calculated and fitted to the experimental data using CRYSOL and aligned on ab initio bead models with SUPCOMB. All the SAXS parameters are summarized in Supplementary Table S3.

### Stopped-flow unwinding assay

Briefly, unwinding kinetics were measured in a two-syringe mode, where *To*Pif1 and fluorescently labeled DNA substrate were pre-incubated at 37 °C in one syringe for 5 min and the unwinding reaction was initiated by rapidly mixing with ATP from another syringe. Each syringe contained unwinding reaction buffer A (25 mM Tris-HCl (pH 7.5), 50 mM NaCl, 2 mM MgCl_2_ and 2 mM DTT). All concentrations listed are after mixing, unless otherwise noted. For converting the output data from volts to percentage unwinding, a calibration experiment was performed in a four-syringe mode, where the helicase, the hexachlorofluorescein-labeled single-stranded oligonucleotides, the fluorescein-labeled single-stranded oligonucleotides and ATP were in four syringes, respectively. The fluorescent signal of the mixed solution from the four syringes corresponded to 100% unwinding. The standard reaction was performed with 4 nM DNA substrates, 1 mM ATP and 100 nM *To*Pif1 in buffer A.

All stopped-flow kinetic traces were averages of ≥10 individual traces. The kinetic traces were analyzed using Bio-Kine (version 4.26, Bio-Logic, France) using Equation 2:

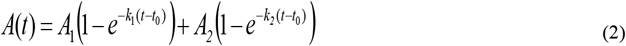

where *A*(t) represents the fraction of DNA unwound at time *t*, *A*_1_ and *A*_2_ are the unwinding amplitudes, *k*_1_ and *k*_2_ are the unwinding rate constants of the two phases, *t*_0_ is the time at which the fraction of DNA unwound starts to rise. From the four parameters obtained through fitting, we can determine the total unwinding amplitude *A*_m_ = *A*_1_ + *A*_2_ and the initial unwinding rate (i.e., the slope of the kinetic unwinding curve at early times) *k*_cat_ = *k*_1_*A*_2_ + *k*_2_*A*_2_.

### Single-molecule fluorescence data acquisition

50 pM fluorescently labeled DNA was added to the chamber containing imaging buffer composed of 20 mM Tris-HCl (pH 7.5), 1 mM MgCl_2_, 50 mM NaCl, 1 mM DTT and an oxygen scavenging system (0.8% D-glucose, 1 mg/mL glucose oxidase, 0.4 mg/mL catalase and 1 mM Trolox). After immobilization for 10 min, free DNA molecules were removed by washing with the imagining buffer. We used an exposure time of 100 ms for all recordings at a constant temperature of 22 °C. FRET efficiency was calculated using *I*_A_/(*I*_D_ + *I*_A_), where *I*_D_ and *I*_A_ represent the intensities of donor and acceptor, respectively. Basic data analysis was carried out by scripts written in MATLAB, and all data fitting were generated using Origin 9.0. Histograms were fitted to Gaussian distributions, with the peak positions unrestrained.

### Circular dichroism spectropolarimetry

Circular dichroism (CD) experiments were performed with a Bio-Logic MOS450/AF-CD optical system (BioLogic Science Instruments, France) equipped with a temperature-controlled cell holder, using a quartz cell with 1-mm path length. A 2.5 μM solution of G4 DNA was prepared in 25 mM Tris–HCl (pH=7.5), 100 mM NaCl. CD spectra were recorded in the UV (220–320 nm) regions in 0.75 nm increments with an averaging time of 2 s at 25°C.

## RESULTS

### ToPif1 unfolds G-quadruplexes in an ATP-dependent manner without topological preference

The DSM specifically targeting parallel G4s is unique feature of the DHX36 helicase family. However, it remains to be determined how G4-unfolding helicases without a specific target motif, such as *To*Pif1, recognize and unfold G4s. To do so, fluorescently labeled parallel and antiparallel intramolecular G4s, 12 nt-ssDNA, and 24 bp blunt-end DNA were titrated with increasing concentrations of *To*Pif1 under equilibrium conditions. The apparent dissociation constant (*K*_d,app_) determined from the titration curves (Figure 1A and Table 1) demonstrated that *To*Pif1 binds both antiparallel (G4^Tel^) and parallel G4s (G4^CEB^) (Supplementary Figure S1A) with similar affinity (*K*_d,app_^antipara^ ≈ 29.63 nM (G4^Tel^) ⩾ *K*_d,app_^para^ ≈ 23.27 nM (G4^CEB^)) without any topological preference. This contrasts sharply with DHX36, which binds parallel G4 with 100-fold higher affinity over nonparallel G4 (17). Furthermore, although both the G4 and ssDNA (T12) were bound with essentially the same affinities (*K*_d,app_^antipara^ ≈ 29.63 nM versus *K*_d,app_^ssDNA^ ≈ 29.12 nM) (Figure 1A (insert) and Table 1), blunt-end DNA bound *To*Pif1 only weakly (*K*_d,app_ > 250.00 nM; Figure 1A (insert) and Table 1). The fact that the apparent dissociation constant of G4 and ssDNA are nearly identical suggests that *To*Pif1 possesses two independent binding sites for ssDNA and G4, respectively. Indeed, competitive binding assays between the labeled G4 and unlabeled ssDNA demonstrated that labeled G4 binding was not influenced by increasing unlabeled ssDNA and vice versa (Supplementary Figure S1B). We then investigated *To*Pif1-mediated G4-unfolding activities by stopped-flow assay according to previous report (35) with both parallel and anti-parallel G4 substrates (36) in the absence or in the presence of ATP. The kinetic data were fit to Equation 2 to determine the unfolding amplitude (*A*_m_) and unfolding rate (*k*_cat_) measured with different G4 DNAs (Figure 1B and Table 1). Although unfolding activity is undetectable in the absence of ATP or in the presence of the non-hydrolysable ATP analog (Supplementary Figure S2), the values of unfolding amplitudes (*A*_m_) and unwinding rates (*k_cat_* (s^−1^)) determined with antiparallel G4s were 1.31-fold and 1.39-fold higher than these determined with parallel G4s (Figure 1B and Table 1).

**Figure 1.**
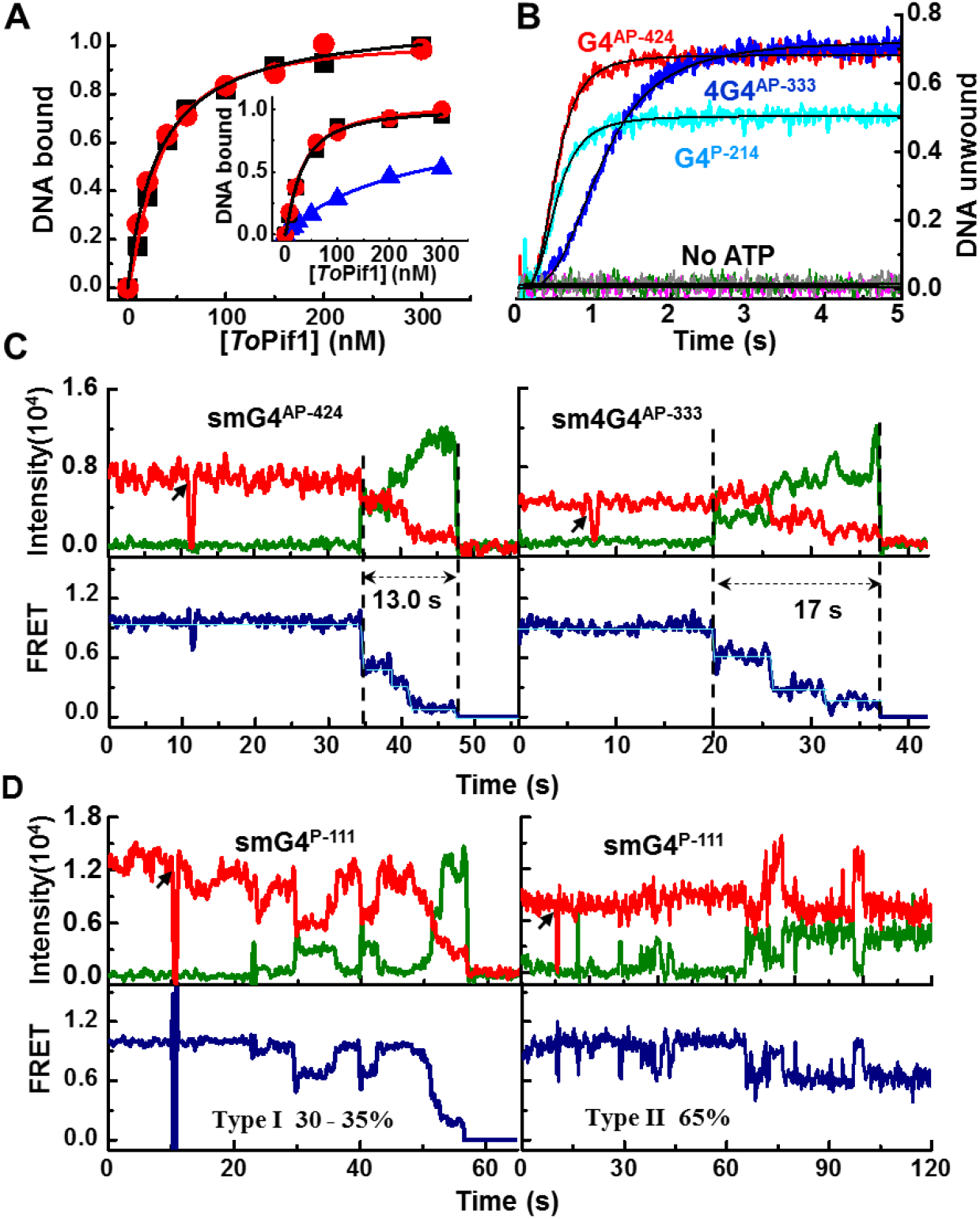
Binding and unwinding activities of *To*Pif1 for different configurations of antiparallel and parallel G4s. **(A)** Comparison of the binding activity of *To*Pif1 for different configurations of antiparallel (G4^Tel^, black points) and parallel (G4^CEB^, red points) G4s. (Insert: Comparison the binding activity of *To*Pif1 to single-stranded DNA (T12, black points) with G4^Tel^, red points), the blue points represent that of the double-strand DNA (D24)). The binding activity was measured using steady-state fluorescence anisotropy assays; 5 nM of fluorescein-labeled DNA substrate and 0.5 mM ADP·AlF_4_ was titrated with increasing protein concentrations at 37 °C. The different binding curves represent an average of 3-4 independent experiments for each substrate. The solid line represents the fit of data according to Eq. (1). **(B)** Stopped-flow DNA unwinding kinetics of *To*Pif1 for different configurations of G4 DNA (G4^AP-424^, 4G4^AP-333^, G4^P-214^) under multiple turnover conditions. All curves represent the average of at least 10 individual traces and the plots are representative of three independent experiments. 4 nM G4 DNA and 100 nM *To*Pif1 were used under experimental conditions as described in ‘Materials and Method’. **(C, D)** The typical smFRET trajectories of antiparallel (smG4^AP-424^, sm4G4^AP-333^) and parallel (smG4^P-111^) G4 unwinding catalyzed by 50 nM *To*Pif1 and 150 μM ATP.

**Table 1.** Binding and unwinding activities of *To*Pif1 for different DNA substrates^1^.

| Substrate |  | Binding | Unwinding |  | 3 |
| --- | --- | --- | --- | --- | --- |
| | | $K_{d,app}$ (nM) | $A_m$ | $k_{cat}$ (s<sup>-1</sup>) | 4 |
| Parallel G4 | G4<sup>CEB</sup> | 23.27 | ND | ND | 5 |
|  | G4<sup>P-214</sup> | ND<sup>2</sup> | 0.55 | 1.57 | 6 |
|  | G4<sup>P-241</sup> | ND | 0.35 | 1.23 | 7 |
|  | G4<sup>P-124</sup> | ND | 0.22 | 1.17 | 8 |
| Antiparallel G4 | G4<sup>Tel</sup> | 29.63 | 0.69 | 1.19 | 9 |
|  | G4<sup>AP-424</sup> | ND | 0.72 | 2.18 | 10 |
|  | 4G4<sup>AP-333</sup> | ND | 0.70 | 0.57 | 11 |
| dsDNA | S<sub>26</sub>D<sub>17</sub> | ND | 0.59 | 3.93 | 12 |
| ssDNA | T<sub>12</sub> | 29.12 | - | - | 13 |
| Blunt-end DNA | D<sub>24</sub> | 250~280 | - | - | 14 |
|  |  |  |  |  | 15 |

<sup>1</sup>Data were determined from 2-3 independent experiments under the experimental conditions as described in Materials and Methods. <sup>2</sup>ND, not determined.

The differences in *To*Pif1-mediated G4 unfolding efficiencies between parallel and antiparallel G4 DNAs were subtle, but significant. To gain mechanistic insight into these differences, smFRET assays were performed with G4 DNAs in which the bases near the 5′ and 3′ sides of G4 were fluorescently labeled and its unfolding activity could be sensitively recorded on smFRET time-traces. The unfolding kinetics of antiparallel G4s comprising three or four G-tetrads (smG4^AP-424^ and sm4G4^AP-333^, respectively) were characterized by clearly stepwise processes in which the smFRET successively decreased from E ≈ 0.90 to E ≈ 0.12 and then finally disappeared due to the escape of the unwound DNA from the coverslip (Figure 1C). The above phenomenon is best interpreted as follows: *To*Pif1 first unfolds G4 from the 5′-lateral G-column, resulting in a G-triplex (G3 with E ≈ 0.58), then further transforms G3 into a G-hairpin structure (E ≈ 0.52) and finally completely unwinds the duplex. Interestingly, for parallel G4 and under the same experimental conditions, although 30-35% time-traces (type I) recorded were essentially the same as those observed with antiparallel G4, ~65% smFRET time-traces (type II) for parallel G4 oscillated between E ≈ 0.90 to E ≈ 0.60 without further decrease before the fluorescence signal bleaching (Figure 1D). Thus, the smFRET assays confirmed that *To*Pif1 unfolds both parallel and antiparallel G4 using the same mechanism, but with a higher efficiency for antiparallel than for parallel G4. The reduced unfolding efficiency in parallel G4 may simply result from the higher thermal stability in the parallel compared with the antiparallel G-quadruplex. Previously reported *T*_m_ values determined with the substrates (G4^AP-424^ and G4^P-214^) showed an inverse correlation with G4 unfolding (36,37). These results are also consistent with previous reports of *Sc*Pif1 unfolding parallel G4 DNA slowly and *Cs*RecQ exclusively unfolding antiparallel G4 DNA (18,24). Altogether, these results may suggest that helicases without DSM display no topological preference for G4 binding and unfolding, and more energy is required to unfold parallel than antiparallel G4 DNA. Furthermore, compared with previous smFRET studies on helicase-mediated G4 unfolding, the smFRET time-trajectory feature of *To*Pif1-mediated antiparallel G4 unfolding is mainly characterized by a stepwise unfolding procedure, without a long oscillation period.

### Overall structure of ToPif1 in complex with G4

To probe the structural basis of *To*Pif1-mediated G4 unfolding, *To*Pif1 was complexed with parallel and antiparallel G4 DNAs flanked at their 5′- and 3′-ends with 6 and 8 nt poly(dT), respectively, thus mimicking G4s formed at a replication fork (38,39). Only parallel G4 (G4^T6/T8^) in complex with *To*Pif1 in a 1:2 ratio in the presence of ADP·AIF_4_ was crystallized and diffracted to 2.58 Å. The asymmetric unit of the crystal structure contained one G4 that was sandwiched between two *To*Pif1 molecules (named molecule *a* and *b*) bound to ssDNAs with 5′→3′ polarity. The two protein molecules in the asymmetric unit were linked through the DNA substrate and related by a rotation of 102.8°, forming dumbbell-sharped structure (Figure 2A and B). Molecule *a* anchored in close proximity to the 5′-most G-tetrad and molecule *b* was bound downstream of the 3′-most G-tetrad. Considering the distance between the binding/unwinding surface constituted by domains 1B/2B and the nearest G-tetrad, in molecule *a* this distance is ≈ 3.00 Å with the 5′-most G-tetrad while in molecule *b* the distance with the last G-tetrad at the 3′ end is greater than 12.00 Å (Figure 2A and B). Therefore, molecule *a* in 5′ is in an active unwinding state during G4 processing, establishing many interactions with G4, while molecule *b* has translocated toward the 3′ end representing a post-catalytic state establishing few contacts with G4 (Figure 2A and B). The domain folding and the spatial arrangement of the modules in *To*Pif1 adopt a similar architecture as those observed in the solved crystal structures of *To*Pif1 (20) and other Pif1 family helicases from yeast (19) and *Bacteroides* species (22). Domains 1A and 2A constitute a deep cleft where ADPoAIF4 is bound and domain 2B is composed of an SH3-like-domain and a prominent β-hairpin (loop3) (Figure 2A and B and Supplementary Figure S3A-D). Each *To*Pif1 molecule binds a 6 nt poly(dT) stretch. Molecule *a* binds T1 to T6 and molecule *b* binds T22 to T27. Conformation of T2-T5 and T24-T27 is conserved and they superpose with a root mean square deviation (RMSD) of 1.20 Å, as already shown for *To*Pif1 complexed with several ligands, ssDNA and ss/dsDNA (20), thus domain 2B exhibits high conformational flexibility. Structural superposition between molecule *a* and the previously determined *To*Pif1 complexed with ss/dsDNA (20) on domain 1A demonstrated that the center of mass of domain 2B moves upwards by 5.40 Å and rotates 25.70° upon G4 binding, and domains 2A/1B, the configurations of bound ssDNA and the amino-acid residues involved in ssDNA binding overlap substantially (RMSD = 0.28 Å over 103 residues) (Supplementary Figure S4A-F), indicating that domain 2B assumes different conformations to accommodate G4 and double-stranded DNA (dsDNA) binding, respectively. Similarly, superposing molecules *a* and *b* on domain 1A shows that G4 binding induced a stable β-hairpin (loop3) formation and the center of mass of domain 2B in molecule *a* moved upwards 5.70 Å, accompanied by 22.90° rotation (Figure 2C), suggesting that domain 2B undergoes a significant conformational adjustment before and after G4 unfolding. G4 is complexed with three K^+^ ions: two are located inside G4 between the tetrads and an extra K^+^ is on top of the 3′ side of G4. This K^+^ ion is stabilized by the last tetrad (tetrad III: G9·G13·G17·G21), T22 and R448 of *To*Pif1 molecule *b*. The overall topology of the G4 structure solved in this work is consistent with these previously reported G4s complexed with proteins, which are characterized by a stack of three G-tetrads and the three short loops containing a single thymine crossing the grooves of the parallel G4 helix joining the top and bottom G-tetrads in the stack (Figure 2D). *To*Pif1-G4^T6/T8^ crystal structure exhibits a good fit with SAXS data (Supplementary Figure S5 and S6 and Table S3) with a χ^2^ of 1.91, confirming that the conformation of observed in the crystal is also the one found in solution.

**Figure 2.**
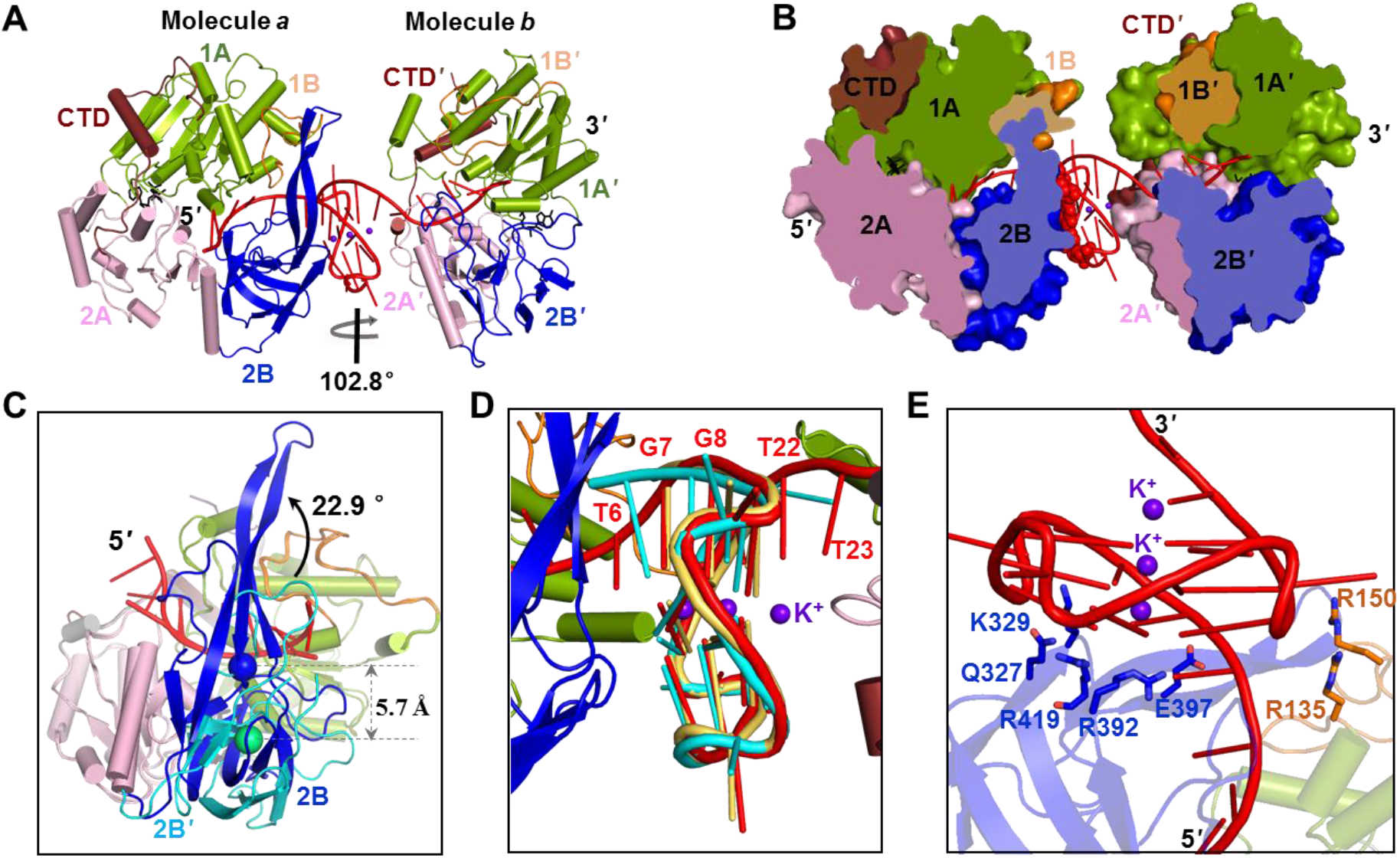
Structure analysis of *To*Pif1 complexed with G-quadruplex (G4) DNA and ADP·AlF_4_. **(A)** Overall structure of *To*Pif1-G4^T6/T8^-ADP·AlF_4_ ternary complex in cartoon mode with the conserved domains 1A (green), 1B (orange), 2A (pink), 2B (blue), and C-terminal (CTD, brown). G4 DNA (G4^T6/T8^) is colored in red and ADP·AlF_4_ shown as a black stick. *To*Pif1 molecules binding at the 5′- and 3′-ends of G4 DNA are labeled as molecule *a* and molecule *b*, respectively. **(B)** Surface view after cutting the crystal structure *To*Pif1-G4^T6/T8^-ADP·AlF_4_ ternary complex and the domains are colored as in (A). **(C)** Structural superposition for the structures of molecules *a* and *b* with the mass centers of domain 2B shown as a sphere. **(D)** Comparison of G4^T6/T8^ (red) in the *To*Pif1 ternary complex, G4^Myc^ (PDB: 5VHE (16)) colored in cyan and G4 ‘T-loops’ (PDB: 6LDM (43)) colored in yellow. **(E)** Residues of molecule *a* that interact with G4 layers in the *To*Pif1-G4^T6/T8^-ADP·AlF4 ternary complex; G4 DNA (G4^T6/T8^) is shown in red and the interacting residues are represented as sticks.

### Structural basis of recognitions and interactions of the entire G-quadruplex and the presumed G-base from the unwound G4

*To*Pif1 recognizes the entire G4 through a cluster of amino acids at domains 1B/2B constituted of G4-Recognizing Surfaces (GRS) (Figure 2E). Although R419 and R392 interact with G11 and G19 in the 5′-most G-tetrad (G7·G11·G15·G19) through cation-π interactions, respectively, the sidechain amino group of Q327 interacts with the phosphate group of G11 through ionic interaction (Figure 3A and Supplementary Figure S7A). Intriguingly, the negatively charged carboxyl group of E397 against the negatively charged oxygen atom on the ribose-phosphate backbone near G7 will exert a repulsion force, which may influence the global conformation of the G4 (Figure 3B). Furthermore, the phosphate group of G20 in the middle tetrad (G8·G12·G16·G20) is stabilized by R135 (Figure 3C). Two positively charged residues, K329 and R150, interact with G9 and G21 in 3′-G-tetrad (G9·G13·G17·G21) through ionic and cation-π interactions (Figure 3D). In sharp contrast to the crystal structure of DHX36 in which just the 5′ hydrophobic G-tetrad is bound by the DSM hydrophobic surface, our structure demonstrated that the residues of GRS form structural clusters which differentially recognize the guanine bases from three different tetrads in the G4. It is therefore reasonable plausible that the spatial conformation of G4 is destabilized by these interactions and G4 is efficiently unfolded in coordination with translocation forces derived from ATP hydrolysis.

**Figure 3.**
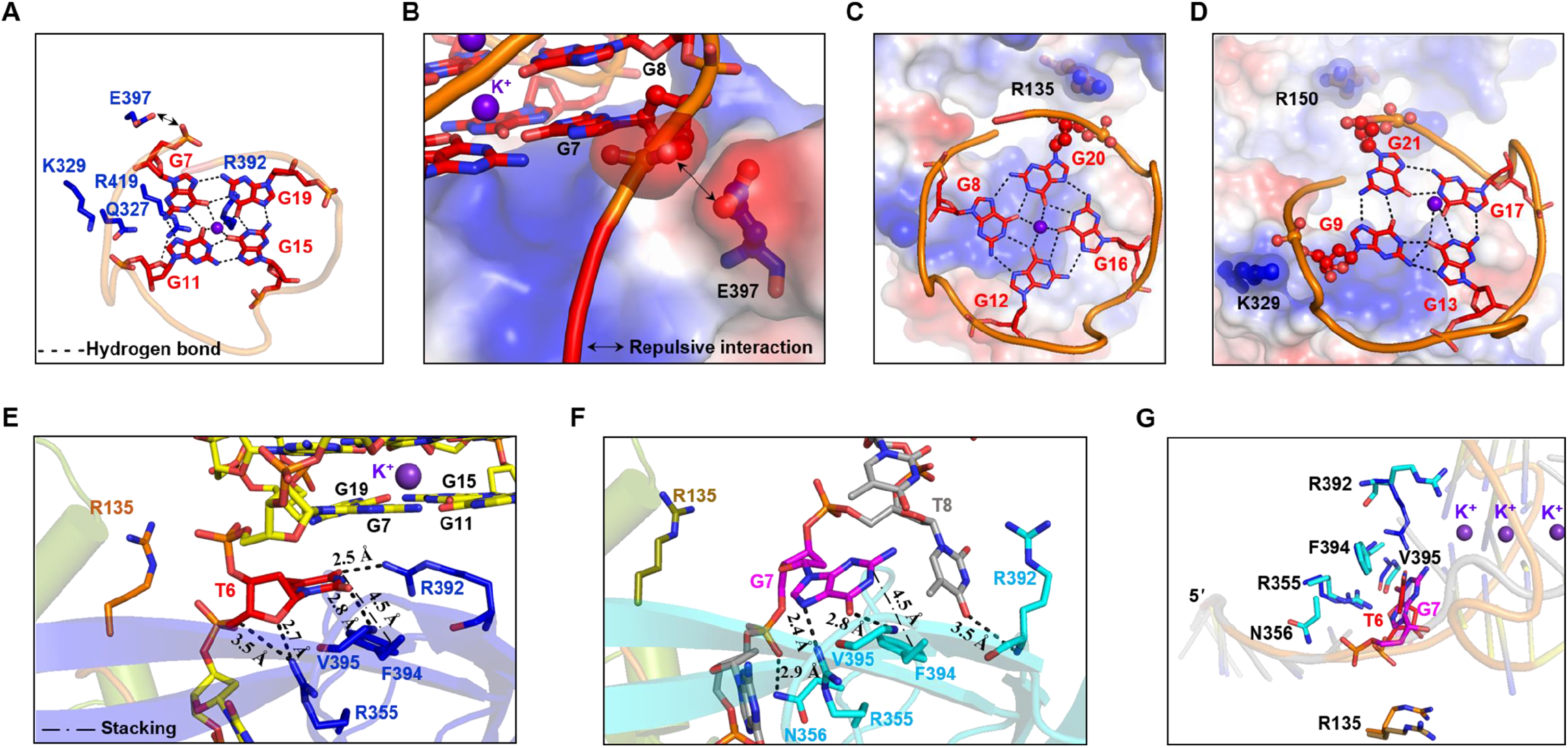
Atomic interactions between the integral G-quadruplex (G4) DNA and *To*Pif1. **(A)** Interactions of *To*Pif1 and the G4 layer I. **(B)**The negative-negative repulsion interaction between E397 and the G7 of G4^T6/T8^ in the *To*Pif1 ternary complex structure. The electrostatic potential at ±5 *kTe^−1^* was colored in blue (basic/positive), white (neutral), and red (acidic/negative). **(C, D)** Interaction of *To*Pif1 and G4 layers II and III, respectively. Molecular electrostatic potential map calculated with G4 DNA omitted from the co-crystal structure (blue to red, ± 5 *kTe^−1^*). **(E)** Interaction between *To*Pif1 and the T6 of G4^T6/T8^ in the *To*Pif1-G4^T6/T8^-ADP·AlF_4_ ternary complex. **(F)**Interaction between *To*Pif1 and the G7 of GR_17_ in the *To*Pif1-GR_17_-ADP·AlF_4_ ternary complex. **(G)** Structural superposition of interacting residues for *To*Pif1 with the T6 of G4^T6/T8^ and the G7 of GR_17_.

In a recently determined crystal structure of *Cs*RecQ in complex with ssDNA bearing a putative G4-forming sequence (human telomere), the 12 nt ssDNA from 3′-end was bound in electropositive channel and the first base (G21) within G-quartet is flipped and sequestrated within just two residues constituting a guanine-specific-pocket (GSP) (18). The conformation of docked G21 in a GSP is interpreted as representing the guanine base from unwound G4, although the potential G4-forming ssDNA sequences were not prefolded to guarantee the formation of a stable G4 in complex with *Cs*RecQ protein. In our *To*Pif1-G4 complex structure (*To*Pif1-G4^T6/T8^), the T6 just upstream of the 5′-tetrad occupying the equivalent position of G21 in *Cs*RecQ (18) is surrounded by a cluster of amino acids and is docked at the compatible GSP position (Figure 3E). T6 also stacks against G19 in the flat 5′-G-tetrad (G7·G11·G15·G19) through a π-π interaction. R392 and V395 are specifically involved in thymine base recognition through hydrogen bonds (R392-T6 = 2.50 Å and V395-T6 = 2.80 Å). Furthermore, the base configuration is further stabilized by π-π stacking between F394 and T6. The residues R355 and R135 interact with the ribose and the phosphodiester of T6, but not its thymine base. To determine whether the binding of a guanine base at the T6 position can restructure the conformation of GRS to accommodate G-base-specific binding, an ssDNA bearing a G4-forming sequence (GR_17_) was directly complexed with *To*Pif1 without prefolding of GR_17_ (20), performed as recently reported to *Cs*RecQ (18). The crystal structure at resolution of 2.21 Å demonstrated that the global conformation of *To*Pif1-GR1_17_ (20) is similar to the complex *To*Pif1-G4^T6/T8^ (Supplementary Figure S7A-C), except that the G4 structure is absent. Structural superposition between the two structures revealed three remarkable features (Figure 3E-G): i) the G7 in binary complex (*To*Pif1-GR1_17_) occupies the equivalent spatial position of T6 in *To*Pif1-G4^T6/T8^; ii) both G7 and T6 flip to the same degree (Figure 3G, ≈113.0°); iii) both the bases and ribose groups of G7 and T6 are bound by the same residues (R392, V395 and R335) with the essentially the same conformation, except that residue N356 establishes an additional hydrogen bond with the phosphate group of G7 upon its binding (Figure 3G). Therefore, if base G7 is considered as the first G base from an unfolded G-tetrad in analogy to G21 in *Cs*RecQ, there is no striking structural reorganization to accommodate the binding of G rather than T. Compared with the previously identified GSP, the cluster of residues surrounding the base (T6/G7) upstream of the G-tetrad in *To*Pif1 are not bound with the equivalent residues identified in the GSP in *Cs*RecQ (Supplementary Figure S8A-C). Of note, the G-rich ssDNAs (GR_17_ (20) and resolved G4 DNA (18)) were not prefolded to form stable G4 DNA in the *To*Pif1 and *Cs*RecQ structures; it therefore remains to be determined whether these structures (*To*Pif1-GR_17_ (20) and CsRecQ-ssDNA (18)) represent an intermediate or/and a product complex of helicases bound to unwound G4 DNA or just ssDNA bound to the proteins.

### Mutational analysis of the residues involved in G4 binding and unfolding

According to the potential functions in G4 binding or/and unfolding, structurally guided single alanine substitution variants of *To*Pif1 were purified to homogeneity, including i) the mutants R392A, R135A and R355A, which interact with the T6/G7 base and/or the ribose/phosphodiester moieties; ii) four variants (Q327A, E397A, R419A and K329A) involved in the 5′-most G-tetrad; and iii) R135 and R150 involved in the interactions with middle G-tetrad and the 3′-most G-tetrad, respectively (Table 2).

**Table 2.** Parameters of binding and unwinding activities of *To*Pif1 and its variants^1^.

| Variants | G-quadruplex |  |  |  | dsDNA |  |  |  |
| --- | --- | --- | --- | --- | --- | --- | --- | --- |
| | $K_{d,app}$ | $A_m$ | $k_{cat}$ | $k_{cat}/K_{d,app}$ | $K_{d,app}$ | $A_m$ | $k_{cat}$ | $k_{cat}/K_{d,app}$ |
| Wild-type<br>*ToPif1* | 29.63 | 0.53 | 0.48 | 0.016 | 30.20 | 0.59 | 3.53 | 0.12 |
| R135A | 119.00 | 0.34 | 0.05 | 0.000 | 30.90 | 0.47 | 1.42 | 0.05 |
| R150A | 152.00 | 0.28 | 0.06 | 0.000 | 31.30 | 0.43 | 2.50 | 0.08 |
| R355A | 133.00 | 0.38 | 0.07 | 0.000 | 38.10 | 0.34 | 1.27 | 0.03 |
| R419A | 38.50 | 0.24 | 0.21 | 0.005 | 38.20 | 0.32 | 2.46 | 0.06 |
| K329A | 48.70 | 0.37 | 0.29 | 0.006 | 30.00 | 0.39 | 3.38 | 0.11 |
| Q327A | 51.50 | 0.48 | 0.41 | 0.008 | 63.40 | 0.43 | 6.14 | 0.10 |
| R392A | 28.50 | 0.40 | 0.27 | 0.009 | 28.30 | 0.32 | 3.50 | 0.12 |
| E397D | 38.80 | 0.45 | 0.35 | 0.010 | 28.00 | 0.49 | 3.27 | 0.12 |
| E397H | 44.60 | 0.78 | 1.32 | 0.030 | 50.90 | 0.31 | 2.64 | 0.05 |
| E397L | 33.00 | 0.88 | 1.18 | 0.036 | 36.00 | 0.48 | 2.36 | 0.06 |
| E397A | 28.30 | 1.00 | 1.19 | 0.042 | 29.40 | 0.49 | 3.79 | 0.13 |

<sup>1</sup>The reported data are the average values determined from 3-4 independent assays as performed under experimental conditions as described in Materials and Methods.

To ascertain qualitatively how the modified residues affect the unfolding activities, the *K*_d,app_ values and the steady-state kinetic parameters, including the unfolding magnitude (*A*_m_) and the unfolding rate (*k*_cat_), were determined with fluorescent-labeled ssDNA and G4 DNA substrates as described above. Several representative G4 unfolding kinetic curves are shown in Figure 4A and the kinetic parameters of all variants are summarized in Table 2. The parameters (*K*_d,app_, *A*_m_ and *k*_cat_) determined with G4 and dsDNA substrates showed a range of variability. To best understand the mutation results, the ratio of *k*_cat_/*K*_d,app_, a useful index for comparing the catalytic efficiency (40), was determined (Table 2). Arranging the G4 *k*_cat_/*K*_d,app_ values in ascending numerical order shows that the relative increase in G4 *k*_cat_/*K*_d,app_ values can be roughly classified into three categories (Figure 4B): i) group I (R355A, R135A and R150A) is characterized by zero values in *K*_cat_/*K*_d,app_ for G4 unfolding while the their *K*_cat_/*K*_d,app_ values determined from dsDNA unwinding are reduced by 33% (R150A) and 75% (R355A); ii) group II (R392A, Q327A, K329A and R419A) is marked by about 44-69% reduction in *k*_cat_/*K*_d,app_ values for G4 unfolding and the corresponding values determined with dsDNA are essentially the same as these determined with wt*To*Pif1, except that *k*_cat_/*K*_d,app_ value determined with R419A for dsDNA is reduced to 50%; iii) group III (E397A, E397Land E397H) displays a surprising results in which *k*_cat_/*K*_d,app_ values determined with G4 are increased to 2-2.6 folds while the *k*_cat_/*K*_d,app_ values for dsDNA are inversely reduced to 50% compared with that of wt*To*Pif. These results will be further analyzed in the next paragraph. Taken together, these results indicate that alteration of the residues in group I-II, particularly the residues in group I, significantly impair G4 unfolding, but only moderately reduce dsDNA unwinding. Consistently, in sharp contrast to wt*To*Pif1, smFRET time trajectories recorded with the mutants in group III revealed that the oscillation curves vary between 0.90 and 0.60, but scarcely attain a completely unfolded level, indicating that the variants can just release the 5’-most lateral G column from G4 DNA, but is unable to unfold the integral G4 DNA completely (Figure 4C-F).

**Figure 4.**
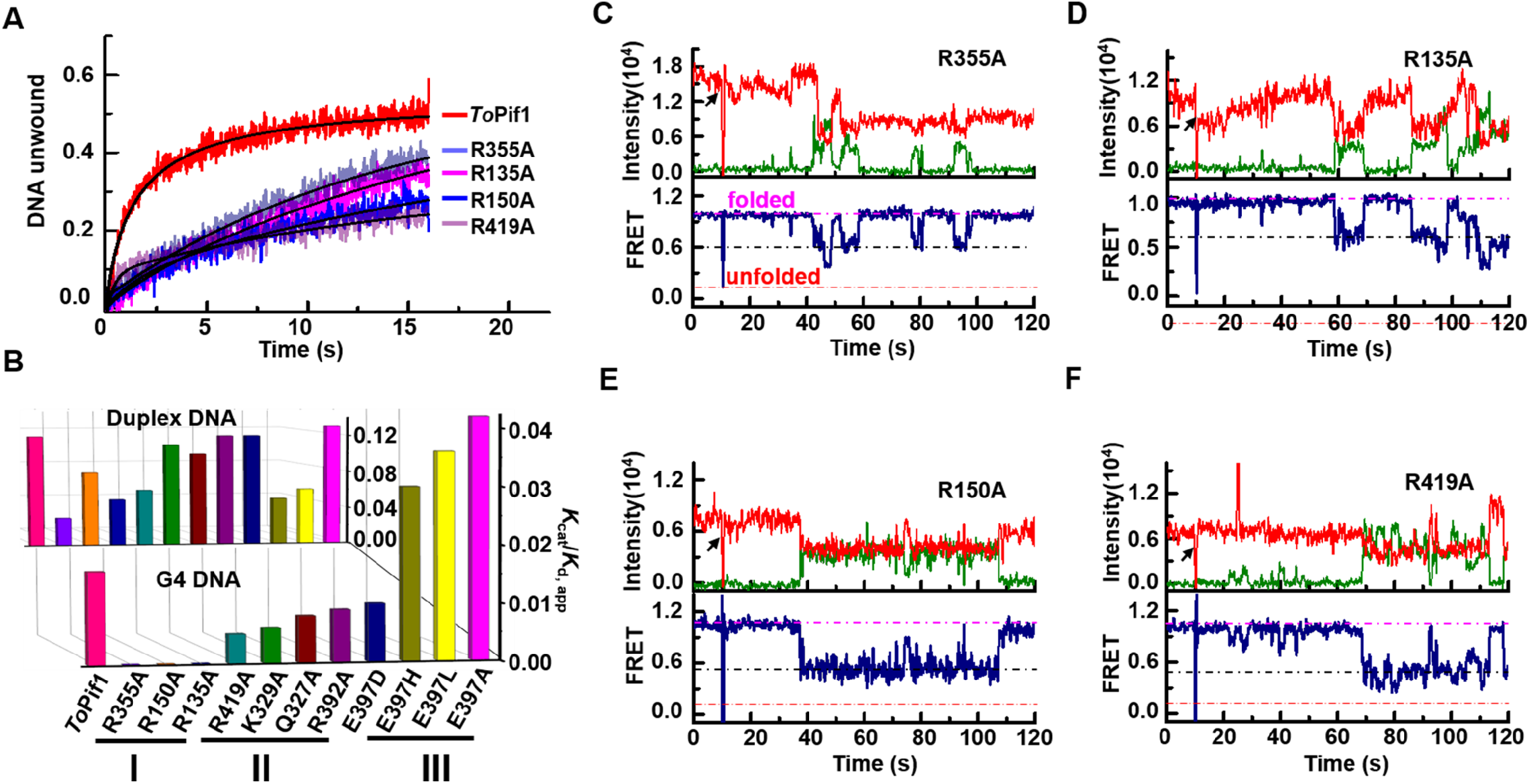
G-quadruplex (G4) unwinding activity of wild-type *To*Pif1 (wt*To*Pif1) and its mutants. **(A)** Stopped-flow kinetics of the G4 (AP-S_16_-TelG4) unwinding activity of wt*To*Pif1 and the various modified proteins. The different unwinding curves represent an average of 3-4 independent experiments for each substrate. The solid line represents the fit of data according to Eq. (2). **(B)** Representation of *k*_cat_/*K*_d,app_ values of G4 in descending numerical order (front panel) and these determined with the dsDNA (S_26_D_17_, back panel). **(C-F)** The typical smFRET trajectories of antiparallel G4 (smG4^AP-424^) unwinding catalyzed by different *To*Pif1 mutants based on single-molecular FRET assays. The pink, black, and red lines represent the states of G4: fully folded, incompletely folded and fully unfolded, respectively. The concentrations of the R355A (C), R135A (D), R150A (E), and R419A (F) mutants were 50 nM with 150 μM ATP. The experimental conditions are described in Methods.

In group III, the kinetic parameters determined with E397A were significantly higher in terms of unfolding amplitude (*A*_m_) and rate (*k*_cat_), respectively (Figure 4B and 5A and Table 2). Compared with wt*To*Pif1, the *k*_cat_/*K*_d,app_ value of E397A determined for G4 unfolding increased by 2.5-fold, although that for dsDNA remained unchanged (Table 2). This stimulation effect was also observed with parallel G4s (Figure 5B). To confirm this interpretation, residue E397 was therefore replaced with a hydrophobic (E397L), a positively charged (E397H) and a negatively charged (E397D) residue. As expected, G4 unfolding activities determined with all variants were systematically higher than those of wt*To*Pif1, except for E397D that — still bearing a negative charge — displayed activity comparable to the wild type (Figure 5A). To gain in-depth mechanistic insight into how residue E397 affects G4-unfolding activity, all of the variants were further studied with smFRET techniques. Analysis of the smFRET time-traces demonstrated that the unfolding times (*t*(*s*)) determined with the three typical variants were significantly reduced, except those for E397D, whose unfolding time remained essentially the same as that of wt*To*Pif1 (Figure 5C-F).

**Figure 5.**
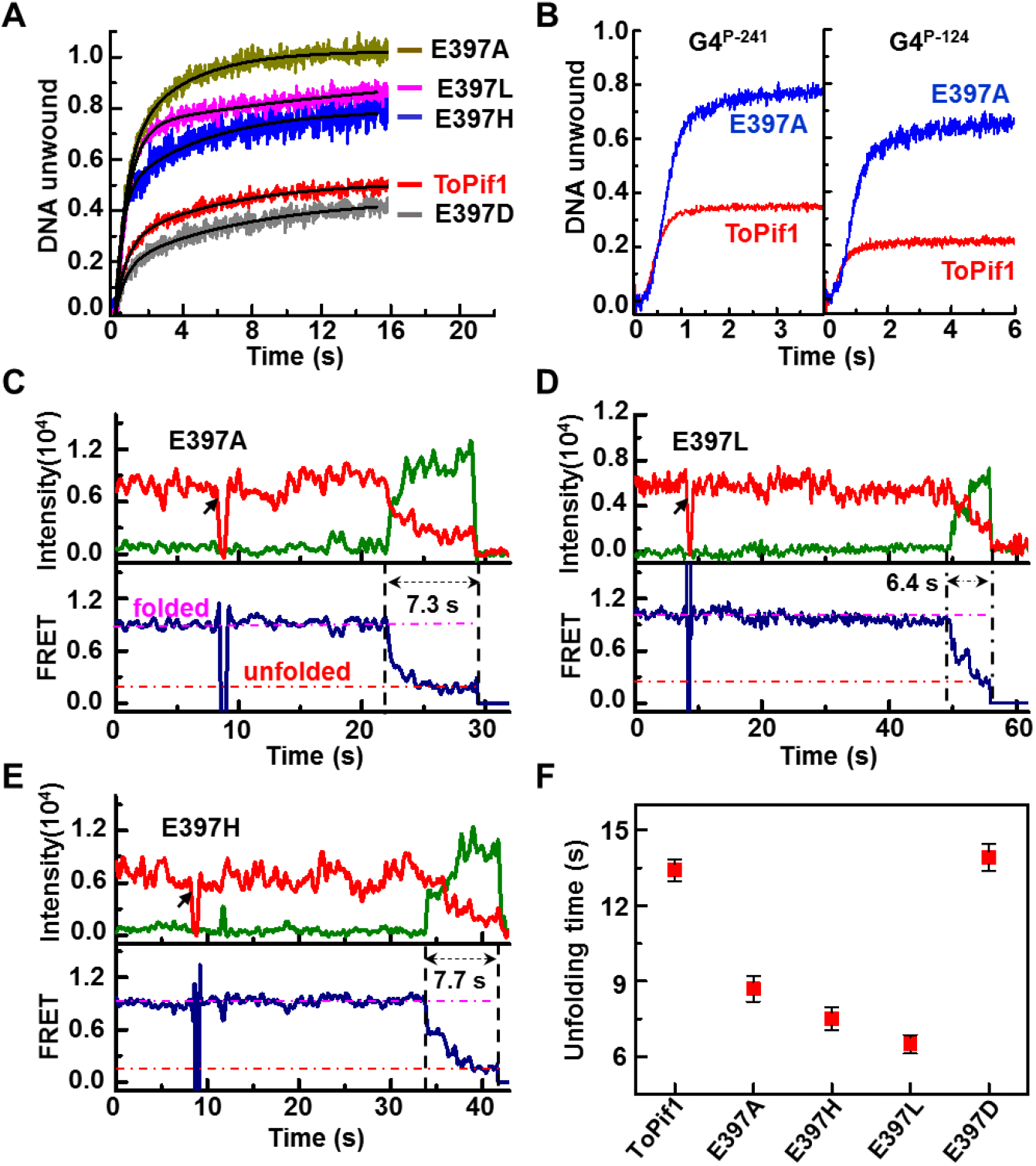
Stimulating effect upon mutation of E397. **(A)** Stopped-flow kinetics of the antiparallel G4 (AP-S_16_-TelG4) unwinding activity of wt*To*Pif1 and its mutants. The different curves represent an average of 3-4 independent experiments for each substrate. The solid line represents the fit of data according to Eq. (2). **(B)** E397A mutant stimulating effect determined with parallel G4 DNA. **(C-E)** The typical FRET trajectories of antiparallel G4 (smG4^AP-424^) unwinding catalyzed by different point mutation on E397 using single-molecular FRET assays. The pink and black lines represent the fully folded and unfolded states of G4, respectively. The different unwinding curves represent an average of 3-4 independent experiments for each substrate. **(F)** Unwinding time of E397A, E397L and E397H determined from unfolding curves of (C), (D), and (E), respectively. The experimental conditions are described in Methods.

The mutation-stimulating G4-resolution activity is surprising and raises the question of why E397A displays higher activity than the wild-type protein and its potential physiological significance. One possibility is that *To*Pif-mediated G4-resolving activity *in vivo* is regulated by an unidentified protein factor. It was recently reported that the proteins Mgs1 and Mms1 in *S. cerevisiae* binding to G4 motifs *in vivo* partially depends on the helicase *Sc*Pif1 (41,42). Although *Sc*Pif1 is the major G4-unwinding helicase in cells, in the absence of Mgs1, the gross chromosomal rearrangement (GCR) rates in yeast is increased, similar to Pif1 deletion (41). It is reasonable to infer two possibilities. First, structural inspection demonstrates that the negatively charged E397 interacts with the phosphate group of G7 within G-tetrad-I and may apply a negative-negative repulsion force, affecting the overall spatial conformation of G4s relative to GRS. Therefore, an unidentified protein in *T. oshimai*, similar to Mgs1 or Mms1 in *S. cerevisiae*, may adjust the spatial conformation of G4 and relieve the negative-negative repulsive force, and consequently stimulate G4-resolving activity. Second, the unidentified protein may interact with *To*Pif1 to activate its G4 unfolding activity. Further studies, based on the work reported here, are needed to complete our understanding of the mechanics of general G4 unfolding mechanisms.

## DISSCUSSION

While increasing evidence has demonstrated that helicase-mediated unfolding of G4s plays an essential role in preserving genome stability, the lack of structural information on G4-processing helicases still hampers our mechanistic understanding of G4 resolution. With a wide array of approaches ranging from bulk/smFRET assays to structural biology analyses, our results reported here provide in-depth insight into the currently largely unknown, but mechanistically important issue of how G4 helicases without a DSM-like motif recognizes the integral G4 structure.

Our binding studies demonstrated that *To*Pif1 possesses strong G4 binding activity without any topological preference. In contrast to the previous reports on DHX36 and the RecQ helicase family, which were reported to use a common mechanism of ATP-independent repetitive unfolding to resolve G4s, *To*Pif1 uses the chemical energy derived from ATP hydrolysis to drive G4 unfolding in a stepwise manner without repetitive cycling. These results imply that *To*Pif1 translocates along the DNA-lattice and sequentially unfolds G4 into G-triplexes and then into G-hairpins. This processing is radically different from that used by DHX36, whose binding triggers the reorganization and destabilization of G4 with a one-at-a-time type of base translocation, and ATP is only needed for releasing the unfolded G4 (16).

DHX36 uses DSM to bind and unfold G4s, most G4-resolving helicases do not possess any equivalent to DSM. An intriguing and unresolved question is how the G4-resolving helicases generally recognize and unfold G4s. We demonstrated that the 1B/2B domains in *To*Pif1 constitute a G4-Recogonizing Surface (GRS) in which the residues recognize the whole G4 structure, not just a tetrad, and differentially interact with the ribose and phosphate units in different tetrads. Superposition of the structural conformation of G4 complexed with *To*Pif1 with that of the DHX36-Myc complex (16) demonstrated that, when the conformation of the G4s are well superposed, loop3 in *To*Pif1 occupies the spatial position of the DSM motif in DHX36, binding on the 5′ side of the G4. Given that Pif1 and DHX36 have opposite polarity for ssDNA translocation, it appears that helicase binding on the 5′ side of G4 is a critical feature for G4 unwinding. We believe that the G4-binding mechanism revealed in this work may be a more general mechanism for helicases. Furthermore, it appears that the previously identified GSP in the RecQ helicase family is not conserved in the *To*Pif1 family, because the residues involved in the first base downstream of 5’-G-tetrad are not selective for guanine and do not form the equivalent hydrogen bonds that stabilize the guanines within G4s as observed in GSPs (Figure 3E-G and Supplementary Figure S8). Finally and interestingly, *To*Pif1-mediated G4-unfolding activity is probably regulated by an unidentified protein which may be similar to Mgs1/Mms1 identified in *S. cerevisiae*, thereby, a whole complex of proteins acts together to control G4 unwinding at different genomic settings. Further studies, instigated by the work reported here, are needed to complete our understanding of the general mechanism underlying how G4-resolving helicases recognize and unfold G4s.

## DATA AVAILABILITY

Accession codes: The atomic coordinate and structure factors have been deposited in the Protein Data Bank (www.wwpdb.org) with the following accession numbers: 7OAR (*To*Pif1-G4^T6/T8^-ADP·AlF_4_).

## SUPPLEMENTARY DATA

Supplementary Data are available at NAR online.

## ACKNOWLEDGEMENTS

We are grateful for access to the SOLEIL (SWING) synchrotron radiation facility for SAXS data collections. We thank the staffs from BL17U1/BL18U1/BL19U1 beamline of National Facility for Protein Science in Shanghai (NFPS) at Shanghai Synchrotron Radiation Facility, for assistance during data collection.

## REFERENCES

1. Burge, S., Parkinson, G.N., Hazel, P., Todd, A.K. and Neidle, S. (2006) Quadruplex DNA: sequence, topology and structure. Nucleic acids research, 34, 5402–5415.

2. Castillo Bosch, P., Segura-Bayona, S., Koole, W., van Heteren, J.T., Dewar, J.M., Tijsterman, M. and Knipscheer, P. (2014) FANCJ promotes DNA synthesis through G-quadruplex structures. Embo j, 33, 2521–2533.

3. Griffin, B.D. and Bass, H.W. (2018) Review: Plant G-quadruplex (G4) motifs in DNA and RNA; abundant, intriguing sequences of unknown function. Plant science : an international journal of experimental plant biology, 269, 143–147.

4. Varshney, D., Spiegel, J., Zyner, K., Tannahill, D. and Balasubramanian, S. (2020) The regulation and functions of DNA and RNA G-quadruplexes. Nature reviews. Molecular cell biology, 21, 459–474.

5. Eddy, J. and Maizels, N. (2006) Gene function correlates with potential for G4 DNA formation in the human genome. Nucleic acids research, 34, 3887–3896.

6. Eddy, J. and Maizels, N. (2008) Conserved elements with potential to form polymorphic G-quadruplex structures in the first intron of human genes. Nucleic acids research, 36, 1321–1333.

7. Huppert, J.L. and Balasubramanian, S. (2005) Prevalence of quadruplexes in the human genome. Nucleic Acids Res, 33, 2908–2916.

8. Todd, A.K., Johnston, M. and Neidle, S. (2005) Highly prevalent putative quadruplex sequence motifs in human DNA. Nucleic acids research, 33, 2901–2907.

9. Schierer, T. and Henderson, E. (1994) A protein from Tetrahymena thermophila that specifically binds parallel-stranded G4-DNA. Biochemistry, 33, 2240–2246.

10. Maizels, N. and Gray, L.T. (2013) The G4 genome. PLoS genetics, 9, e1003468.

11. Fontana, G.A. and Gahlon, H.L. (2020) Mechanisms of replication and repair in mitochondrial DNA deletion formation. Nucleic acids research, 48, 11244–11258.

12. De, S. and Michor, F. (2011) DNA secondary structures and epigenetic determinants of cancer genome evolution. Nature structural & molecular biology, 18, 950–955.

13. Edwards, D.N., Machwe, A., Wang, Z. and Orren, D.K. (2014) Intramolecular telomeric G-quadruplexes dramatically inhibit DNA synthesis by replicative and translesion polymerases, revealing their potential to lead to genetic change. PloS one, 9, e80664.

14. Eddy, S., Maddukuri, L., Ketkar, A., Zafar, M.K., Henninger, E.E., Pursell, Z.F. and Eoff, R.L. (2015) Evidence for the kinetic partitioning of polymerase activity on G-quadruplex DNA. Biochemistry, 54, 3218–3230.

15. Mendoza, O., Bourdoncle, A., Boulé, J.-B., Brosh, R.M., Jr and Mergny, J.-L. (2016) G-quadruplexes and helicases. Nucleic acids research, 44, 1989–2006.

16. Chen, M.C., Tippana, R., Demeshkina, N.A., Murat, P., Balasubramanian, S., Myong, S. and Ferré-D’Amaré, A.R. (2018) Structural basis of G-quadruplex unfolding by the DEAH/RHA helicase DHX36. Nature, 558, 465–469.

17. Heddi, B., Cheong, V.V., Martadinata, H. and Phan, A.T. (2015) Insights into G-quadruplex specific recognition by the DEAH-box helicase RHAU: Solution structure of a peptide-quadruplex complex. Proc Natl Acad Sci U S A, 112, 9608–9613.

18. Voter, A.F., Qiu, Y., Tippana, R., Myong, S. and Keck, J.L. (2018) A guanine-flipping and sequestration mechanism for G-quadruplex unwinding by RecQ helicases. Nature communications, 9, 4201.

19. Lu, K.Y., Chen, W.F., Rety, S., Liu, N.N., Wu, W.Q., Dai, Y.X., Li, D., Ma, H.Y., Dou, S.X. and Xi, X.G. (2018) Insights into the structural and mechanistic basis of multifunctional S. cerevisiae Pif1p helicase. Nucleic acids research, 46, 1486–1500.

20. Dai, Y.X., Chen, W.F., Liu, N.N., Teng, F.Y., Guo, H.L., Hou, X.M., Dou, S.X., Rety, S. and Xi, X.G. (2021) Structural and functional studies of SF1B Pif1 from Thermus oshimai reveal dimerization-induced helicase inhibition. Nucleic acids research, 49, 4129–4143.

21. Dehghani-Tafti, S., Levdikov, V., Antson, A.A., Bax, B. and Sanders, C.M. (2019) Structural and functional analysis of the nucleotide and DNA binding activities of the human PIF1 helicase. Nucleic acids research, 47, 3208–3222.

22. Zhou, X., Ren, W., Bharath, S.R., Tang, X., He, Y., Chen, C., Liu, Z., Li, D. and Song, H. (2016) Structural and Functional Insights into the Unwinding Mechanism of Bacteroides sp Pif1. Cell Rep, 14, 2030–2039.

23. Li, J.H., Lin, W.X., Zhang, B., Nong, D.G., Ju, H.P., Ma, J.B., Xu, C.H., Ye, F.F., Xi, X.G., Li, M. et al. (2016) Pif1 is a force-regulated helicase. Nucleic acids research, 44, 4330–4339.

24. Wang, L., Wang, Q.M., Wang, Y.R., Xi, X.G. and Hou, X.M. (2018) DNA-unwinding activity of Saccharomyces cerevisiae Pif1 is modulated by thermal stability, folding conformation, and loop lengths of G-quadruplex DNA. J Biol Chem, 293, 18504–18513.

25. Dahan, D., Tsirkas, I., Dovrat, D., Sparks, M.A., Singh, S.P., Galletto, R. and Aharoni, A. (2018) Pif1 is essential for efficient replisome progression through lagging strand G-quadruplex DNA secondary structures. Nucleic acids research, 46, 11847–11857.

26. Li, S.B., Wang, H.L., Jehi, S.A., Li, J., Liu, S., Wang, Z., Truong, L., Chiba, T., Wang, Z.F. and Wu, X.H. (2021) PIF1 helicase promotes break-induced replication in mammalian cells. Embo J, 40.

27. Zhang, W.-Z., Tang, J.-C., Wang, S.-S., Wang, Z.-J., Qin, W.-M. and He, J.-H. (2019) The protein complex crystallography beamline (BL19U1) at the Shanghai Synchrotron Radiation Facility. Nuclear Science and Techniques, 30, 170.

28. Kabsch, W. (2010) XDS. Acta Crystallogr D Biol Crystallogr, 66, 125–132.

29. Adams, P.D., Afonine, P.V., Bunkoczi, G., Chen, V.B., Davis, I.W., Echols, N., Headd, J.J., Hung, L.W., Kapral, G.J., Grosse-Kunstleve, R.W. et al. (2010) PHENIX: a comprehensive Python-based system for macromolecular structure solution. Acta crystallographica. Section D, Biological crystallography, 66, 213–221.

30. McCoy, A.J., Grosse-Kunstleve, R.W., Adams, P.D., Winn, M.D., Storoni, L.C. and Read, R.J. (2007) Phaser crystallographic software. J Appl Crystallogr, 40, 658–674.

31. Lovelace, L.L., Minor, W. and Lebioda, L. (2005) Structure of human thymidylate synthase under low-salt conditions. Acta Crystallogr D Biol Crystallogr, 61, 622–627.

32. Emsley, P., Lohkamp, B., Scott, W.G. and Cowtan, K. (2010) Features and development of Coot. Acta Crystallogr D Biol Crystallogr, 66, 486–501.

33. David, G. and Pérez, J. (2009) Combined sampler robot and high - performance liquid chromatography: a fully automated system for biological small - angle X - ray scattering experiments at the Synchrotron SOLEIL SWING beamline. Journal of Applied Crystallography, 42, 892–900.

34. Petoukhov, M.V., Franke, D., Shkumatov, A.V., Tria, G., Kikhney, A.G., Gajda, M., Gorba, C., Mertens, H.D., Konarev, P.V. and Svergun, D.I. (2012) New developments in the ATSAS program package for small-angle scattering data analysis. J Appl Crystallogr, 45, 342–350.

35. Liu, N.N., Ji, L., Guo, Q., Dai, Y.X., Wu, W.Q., Guo, H.L., Lu, K.Y., Li, X.M. and Xi, X.G. (2020) Quantitative and real-time measurement of helicase-mediated intra-stranded G4 unfolding in bulk fluorescence stopped-flow assays. Analytical and bioanalytical chemistry, 412, 7395–7404.

36. Cheng, M., Cheng, Y., Hao, J., Jia, G., Zhou, J., Mergny, J.L. and Li, C. (2018) Loop permutation affects the topology and stability of G-quadruplexes. Nucleic acids research, 46, 9264–9275.

37. Bugaut, A. and Balasubramanian, S. (2008) A sequence-independent study of the influence of short loop lengths on the stability and topology of intramolecular DNA G-quadruplexes. Biochemistry, 47, 689–697.

38. Lee, W.T.C., Yin, Y., Morten, M.J., Tonzi, P., Gwo, P.P., Odermatt, D.C., Modesti, M., Cantor, S.B., Gari, K., Huang, T.T. et al. (2021) Single-molecule imaging reveals replication fork coupled formation of G-quadruplex structures hinders local replication stress signaling. Nature Communications, 12, 2525.

39. Lemmens, B., van Schendel, R. and Tijsterman, M. (2015) Mutagenic consequences of a single G-quadruplex demonstrate mitotic inheritance of DNA replication fork barriers. Nat Commun, 6, 8909.

40. Barlow, J.N., Conrath, K. and Steyaert, J. (2009) Substrate-dependent modulation of enzyme activity by allosteric effector antibodies. Biochimica et biophysica acta, 1794, 1259–1268.

41. Paeschke, K. and Burkovics, P. (2021) Mgs1 function at G-quadruplex structures during DNA replication. Current genetics, 67, 225–230.

42. Schwindt, E. and Paeschke, K. (2018) Mms1 is an assistant for regulating G-quadruplex DNA structures. Current genetics, 64, 535–540.

43. Traczyk, A., Liew, C.W., Gill, D.J. and Rhodes, D. (2020) Structural basis of G-quadruplex DNA recognition by the yeast telomeric protein Rap1. Nucleic acids research, 48, 4562–4571.

