## supplementary information for "Structural mechanism underpinning *Thermus oshimai* Pif1-mediated G-quadruplex unfolding"

*Footnotes:*

[†]These authors contributed equally to this work.

**This PDF file includes:**

Supplementary Tables S1-S3

Supplementary Figures S1-S8

References for SI reference citations

### SUPPLEMENTARY TABLES

**Supplementary Table S1.** DNA sequences used in assays.

| Assay | Name | Sequence |
|---|---|---|
| Binding | G4$^{Tel}$ | 5' GGGTTAGGGTTAGGGTTAGGG-Fam |
| | G4$^{CEB}$ | 5' AGGGTGGGTGGGTGGG-Fam |
| | $T_{12}$ | 5' TTTTTTTTTTTT-Fam |
| | $D_{24}$ | 5' GCCCTGGTGCCGACCAACGAAGGT-Fam<br>3' CGGGACCACGGCTGGTTGCTTCCA |
| DNA unwinding (stopped-flow) | G4$^{AP\text{-}424}$ | 5' GTGTTGATGAAGGGTTTTGGGTTGGGTTTTGGGAGGATCCTCGAG-Fam |
| | 4G4$^{AP\text{-}333}$ | 5' GTGTTGATGAAGGGGTTAGGGGTTAGGGGTTAGGGGAGGATCCTCGAG-Fam |
| | G4$^{P\text{-}214}$ | 5' GTGTTGATGAAGGGTTGGGTGGGTTTTGGGAGGATCCTCGAG-Fam |
| | G4$^{P\text{-}241}$ | 5' GTGTTGATGAAGGGTTGGGTTTTGGGTGGGAGGATCCTCGA*G*-Fam |
| | G4$^{P\text{-}124}$ | 5' GTGTTGATGAAGGGTGGGTTGGGTTTTGGGAGGATCCTCGAG-Fam |
| | 5HF-G4 parten | 5' Hex-CTCGAGGATCCTTT |
| | AP-S$_{16}$-TelG4 | 5' (T)$_{16}$GGGT$_{(ATTO)}$TAGGGTTAGGGTTAGGGA$_{(Cy3)}$ |
| | S$_{26}$D$_{17}$ | 5' (T)$_{26}$ATGTATGTCAAGGAAGG-Fam<br>3' TACATACAGTTCCTTCC-Hex |
| smFRET | smG4$^{AP\text{-}424}$ | 5' TGTGAGCGTGGT$_{(Cy3)}$GTGGGTTTTGGGTTGGGTTTTGGGATGTATGACAAGGAAGG |
| | smG4$^{P\text{-}111}$ | 5' TGTGAGCGTGGT$_{(Cy3)}$GTGGGAGGGAGGGAGGGATGTATGACAAGGAAGG |
| | sm4G4$^{AP\text{-}333}$ | 5' TGTGAGCGTGGT$_{(Cy3)}$GTGGGGTTAGGGGTTAGGGGTTAGGGGATGTATGACAAGGAAGG |
| | 17nt-biotin | 3' T$_{(Cy5)}$ACATACTGTTCCTTCC-Biotin |
| Crystallization | G4$^{T6/T8}$ | 5' (T)$_{6}$GGGTGGGTGGGTGGG(T)$_{8}$ |
| | GR$_{17}$ | 5' GGTTTGGTTTGGTTTGG |

**Supplementary Table S2.** Crystallization data collection and refinement statistics.[a]

| | ***To*Pif1-G4$^{T6/T8}$-ADP·AlF$_4$** |
|---|---|
| **PDB ID code** | 7OAR |
| **Data collection** | |
| Wavelength (Å) | 0.9791 |
| Resolution range (Å) | 32.86 - 2.58 (2.67 - 2.58) |
| Space group | $P6_2\ 2\ 2$ |
| Unit cell | |
| *a*, *b*, *c* (Å) | 151.78 151.78 219.53 |
| Multiplicity | 19.9 (20.5) |
| Completeness (%) | 94.1 (46.0) |
| Mean I/sigma (I) | 26.8 (1.4) |
| R-merge | 0.069 (2.402) |
| CC1/2 | 0.997 (0.651) |
| **Refinement** | |
| Reflections used in refinement | 44511 (1876) |
| R-work | 0.199 (0.327) |
| R-free | 0.245 (0.360) |
| Number of atoms | |
| Macromolecule | 7421 |
| Ligand | 69 |
| Solvent | 38 |
| RMS (bonds) | 0.008 |
| RMS (angles) | 1.16 |
| Ramachandran favored (%) | 99.76 |
| Ramachandran allowed (%) | 0.24 |
| Average B-factor | 113.67 |
| Macromolecule | 111.65 |
| Ligand | 89.61 |
| Solvent | 82.78 |

[a] Statistics for the highest-resolution shell are shown in parentheses.

**Supplementary Table S3.** SAXS data collection and parameters.

| | **ToPif1+G4$^{T6/T8}$** |
|---|---|
| **Data-collection parameters** | |
| Instrument | SWING |
| Beam geometry (mm) | 0.4×0.1 |
| Wavelength (Å) | 1.03 |
| q range (Å$^{-1}$) | 0.007-0.5 |
| Detector | Aviex |
| Data collection mode | HPLC |
| Exposure time (s) / nb frames | 1 / 255 |
| Concentration (injected) (mg.ml$^{-1}$) | 10 |
| Temperature (K) | 288 |
| **Structural parameters** | |
| Guinier quality: | |
| data points | 3-54 |
| *qR*g | 0.24-1.21 |
| correlation coefficient | 0.993 |
| *I*(0) (cm$^{-1}$) [from Guinier] | 5.199 ± 0.008 |
| *R*g (Å) [from Guinier] | 36.67 ± 0.06 |
| *R*g (Å) [from *P*(*r*)] | 36.41 |
| $D_{max}$ (Å) | 120.0 |
| Porod estimate (Å$^3$) | 156394 |
| **Molecular-mass determination** | |
| Partial specific volume (cm$^3$.g$^{-1}$) | 0.64 |
| Contrast ($\Delta\rho \times 10^{10}$ cm$^{-2}$) | 4.510 |
| Molecular mass $M_r$ [from *I*(0)] | 121800 |
| Calculated monomeric $M_r$ from sequence | 57394 + 57394 + 9138=123926 |
| **Data processing** | |
| Primary data reduction | FOXTROT |
| Data processing | PRIMUS |
| *Ab initio* analysis | DAMMIF |
| Number of models | 50 |
| Model $\chi^2$ | 2.61± 0.03 |
| Validation and averaging | DAMAVER |
| NSD | 0.79 ± 0.25 |
| Estimated resolution (Å) | 46 ± 3 |
| Rigid-body modelling | SASREF |
| Computation of model intensities | CRYSOL |
| Model $\chi^2$ | 1.905 |

### SUPPLEMENTARY FIGURES

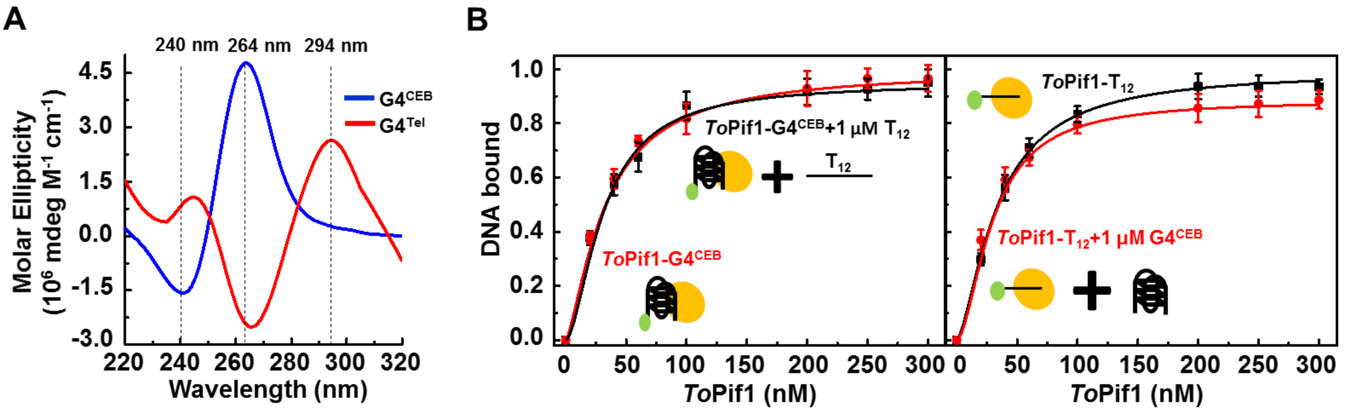


**Supplementary Figure S1. Competitive binding assays between G4 and ssDNA. (A)** CD experiments of G4 DNA. **(B)** Left panel: DNA binding of *To*Pif1 to $G4^{CEB}$ with (black line) or without (red line) 1 μM ssDNA ($T_{12}$). Fitting of the data to the Eq.1 yields a $K_{d,app}$ value for *To*Pif1-$G4^{CEB}$ of 30.30 ± 1.17 nM, a $K_{d,app}$ value for *To*Pif1-$G4^{CEB}$ (added with 1 μM $T_{12}$) of 30.91 ± 2.32 nM. Right panel: DNA binding of *To*Pif1 to $T_{12}$ with (red line) or without (black line) 1 μM G4 DNA ($G4^{CEB}$). Fitting of the data to the Eq.1 yields a $K_{d,app}$ value for *To*Pif1-$T_{12}$ of 29.10 ± 0.53 nM, a $K_{d,app}$ value for *To*Pif1-$T_{12}$ (added with 1 μM $G4^{CEB}$) of 26.50 ± 0.71 nM. DNA binding assays were carried out in buffer A (25 mM Tris-HCl (pH 7.5), 50 mM NaCl, 2 mM $MgCl_2$ and 2 mM DTT) with 0.5 mM ADP·$AlF_4$.

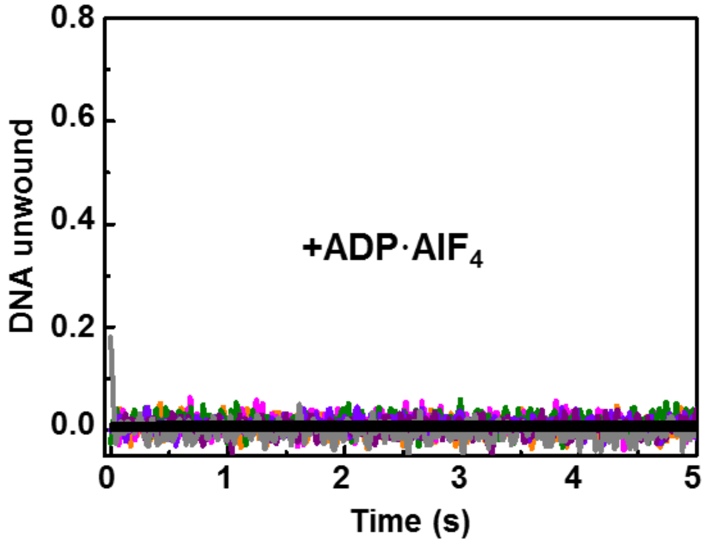


**Supplementary Figure S2. Unfolding activities of *To*Pif1 in the presence of non-hydrolysable ATP analogs.** Stopped-flow unwinding kinetic curves of *To*Pif1 unwinding G4 DNA ($G4^{AP\text{-}424}$, $G4^{P\text{-}214}$, $4G4^{AP\text{-}333}$) in buffer A with 1 mM ADP·$AlF_4$.

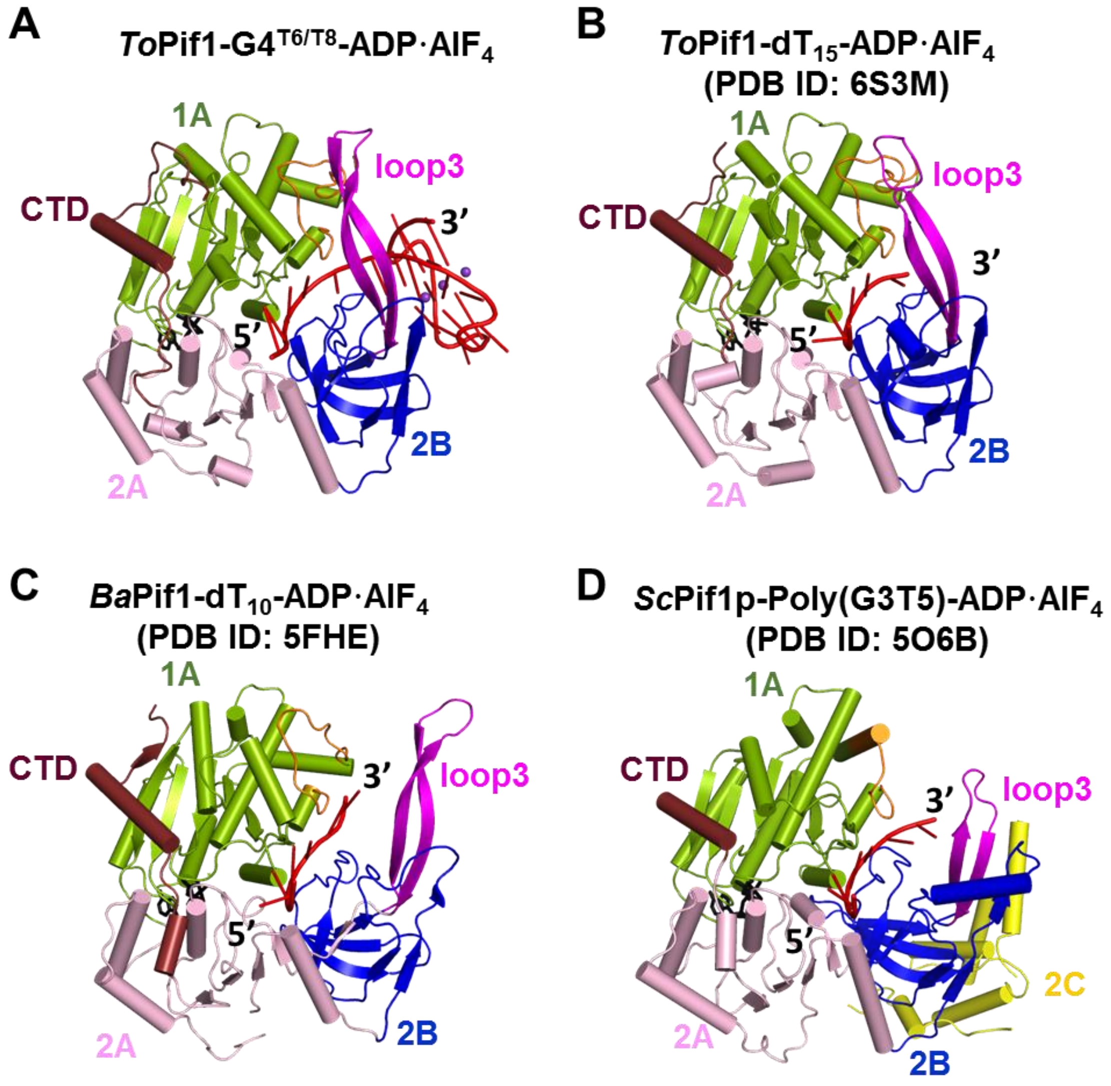


**Supplementary Figure S3. Domain folding and spatial arrangement of *To*Pif1 in a ternary complex (*To*Pif1-G4$^{T6/T8}$-ADP·AlF$_4$) adopt an architecture similar to that of previously solved Pif1 helicase structures. (A)** Cartoon representation of *To*Pif1 (molecule *a*) with G4$^{T6/T8}$ in the presence of ADP·AlF$_4$ shown as black stick and loop3 highlighted in magenta. **(B)** Cartoon representation of *To*Pif1-dT$_{15}$-ADP·AlF$_4$ (PDB: 6S3M (1)) with ADP·AlF$_4$ shown as black stick and loop3 highlighted in magenta. **(C)** Cartoon representation of BaPif1-dT$_{10}$- ADP·AlF$_4$ (PDB: 5FHE (2)) with ADP·AlF$_4$ shown as black stick and loop3 highlighted in magenta. **(D)** Cartoon representation of *Sc*Pif1p-poly(G3T5)-ADP·AlF$_4$ (PDB: 5O6B (3)) with ADP·AlF$_4$ shown as black stick and loop3 highlighted in magenta.

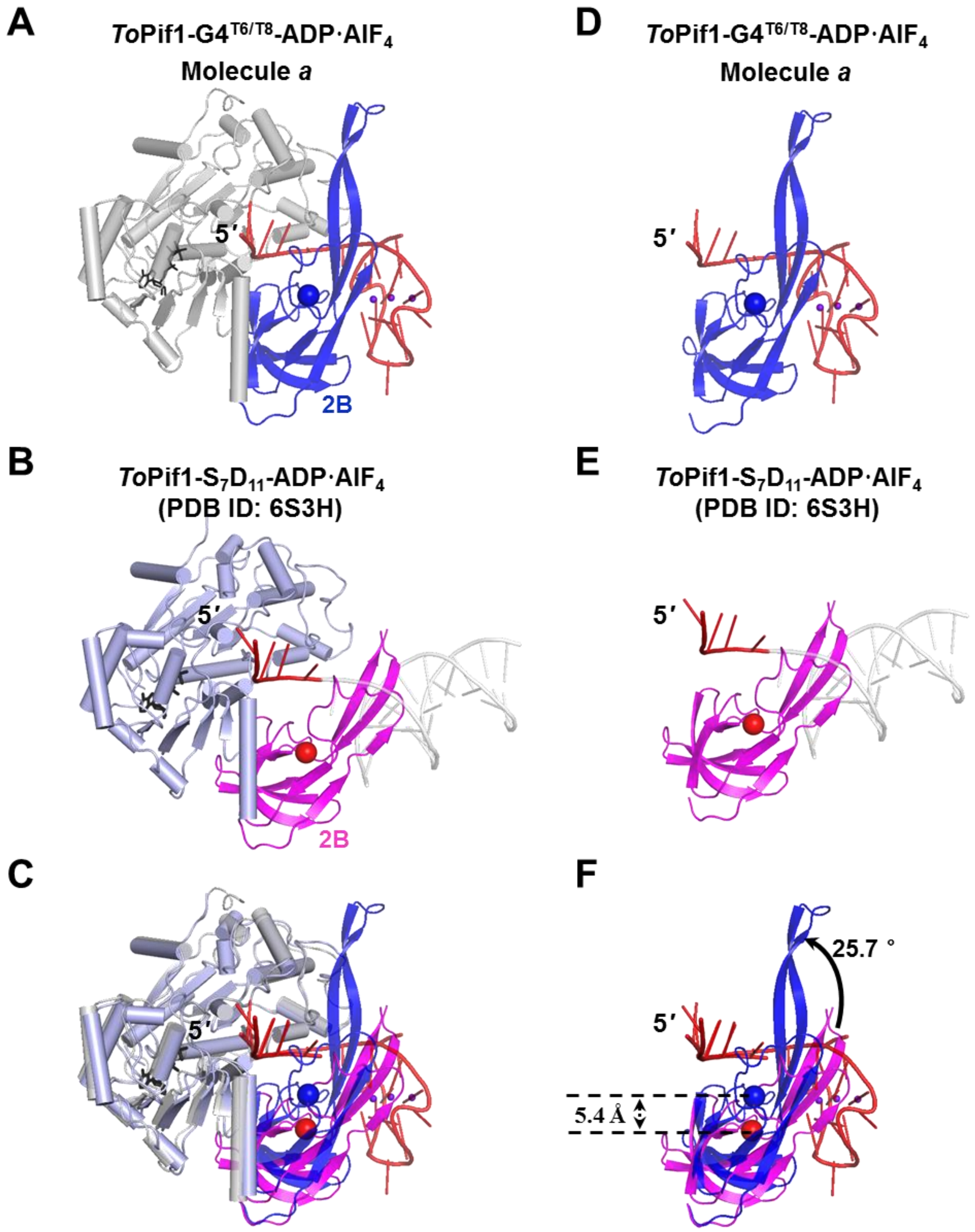


**Supplementary Figure S4. Comparison of domain 2B of molecule *a* of the *To*Pif1-G4$^{T6/T8}$-ADP·AlF$_4$ ternary complex and the previously determined *To*Pif1-ss/dsDNA structure (PDB: 6S3H (1)). (A)** Cartoon representation of molecule *a* in the *To*Pif1-G4$^{T6/T8}$-ADP·AlF$_4$ structure with domain 2B and the center mass (sphere) of domain 2B highlighted in blue. G-quadruplex (G4) DNA is shown in red. **(B)** Cartoon representation of the *To*Pif1-S$_7$D$_{11}$-ADP·AlF$_4$ structure (PDB: 6S3H (1)) with domain 2B shown in magenta. The center mass of domain 2B is shown as a red sphere. **(C)** Superposition on domain 1A of the structures in (A) and (B). **(D)** Domain 2B and G4 DNA of molecule *a* in the *To*Pif1-G4$^{T6/T8}$-ADP·AlF$_4$ structure. **(E)** Domain 2B and modeled duplex DNA in the *To*Pif1-S$_7$D$_{11}$-ADP·AlF$_4$ structure (PDB: 6S3H (1)). **(F)** Superposition on domain 1A of the structure in (C) with only domain 2B and G4 DNA being shown.

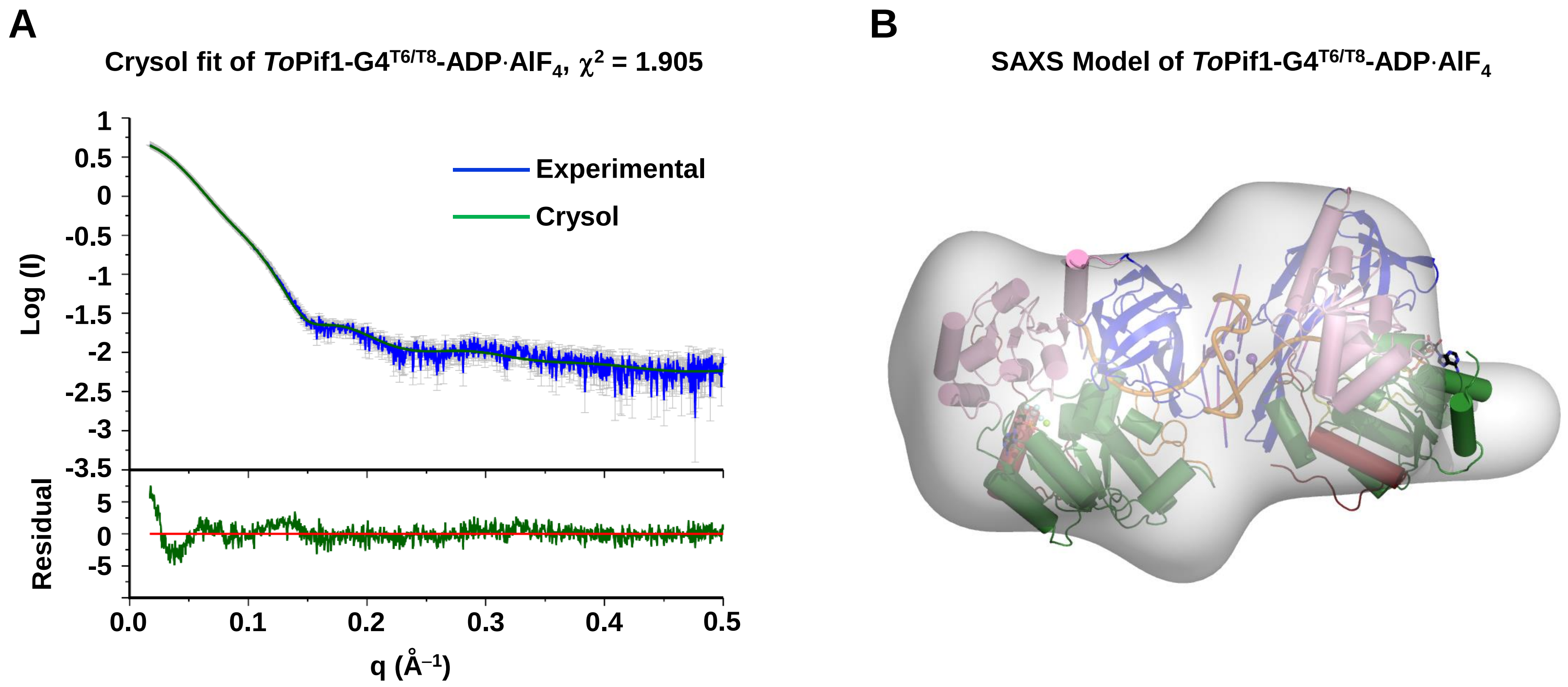


**Supplementary Figure S5. SAXS result of *To*Pif1-G4$^{T6/T8}$-ADP·AlF$_4$. (A)** Fit curve of the SAXS data of *To*Pif1- G4$^{T6/T8}$-ADP·AlF$_4$ with two modelled *To*Pif1 molecules bound to the substrate G4$^{T6/T8}$ calculated with Crysol. **(B)** The model of *To*Pif1- G4$^{T6/T8}$-ADP·AlF$_4$ superimposed on the *ab initio* envelope calculated with DAMMIF.

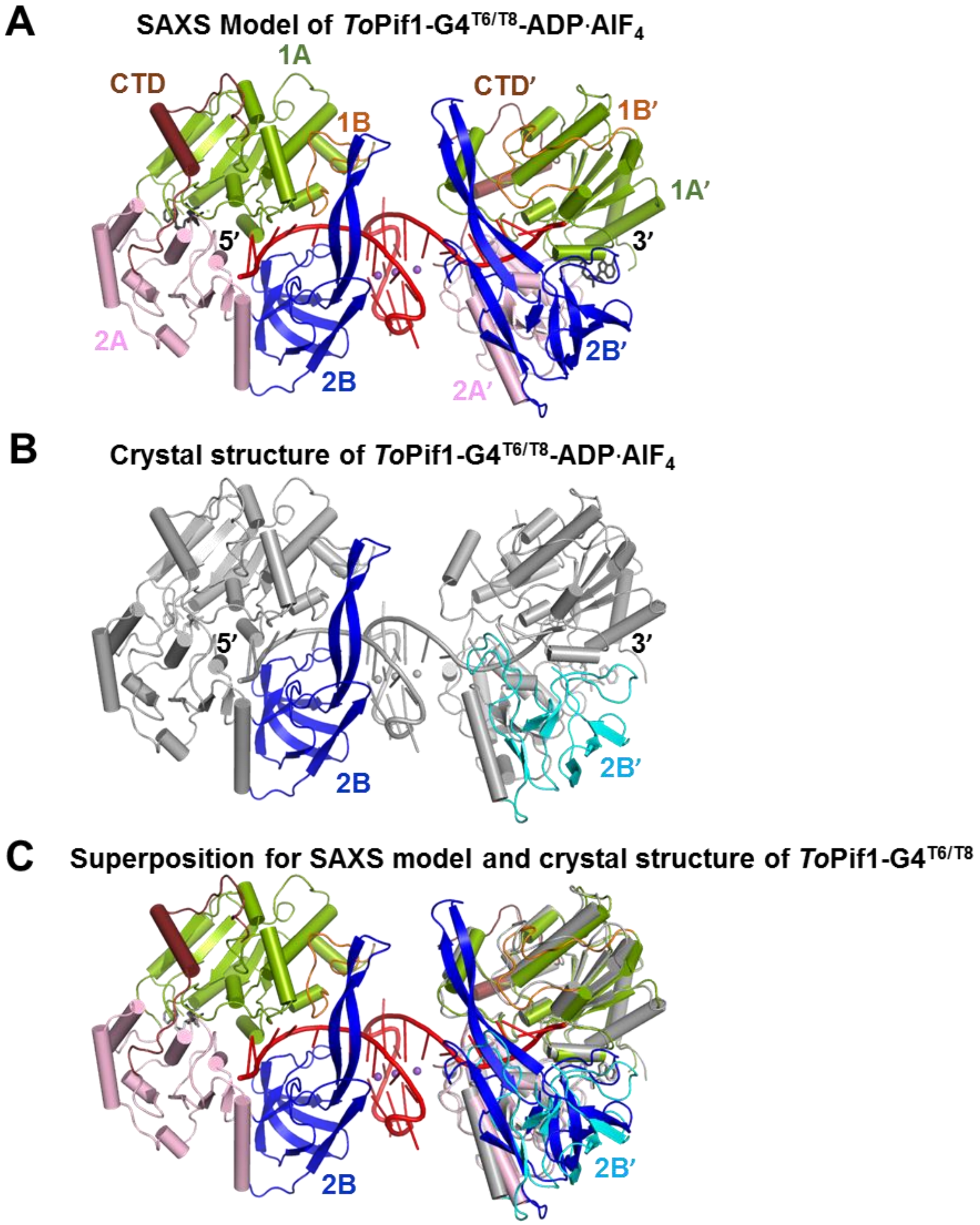


**Supplementary Figure S6. Comparison for SAXS model and crystal structure of *To*Pif1-G4$^{T6/T8}$-ADP·AlF$_{4.}$ (A)** The SAXS Model of *To*Pif1-G4$^{T6/T8}$ -ADP·AlF$_4$. **(B)** Crystal structure of *To*Pif1-G4$^{T6/T8}$ -ADP·AlF$_{4.}$ **(C)** Superposition for SAXS model and crystal structure of *To*Pif1-G4$^{T6/T8}$ -ADP·AlF$_4$.

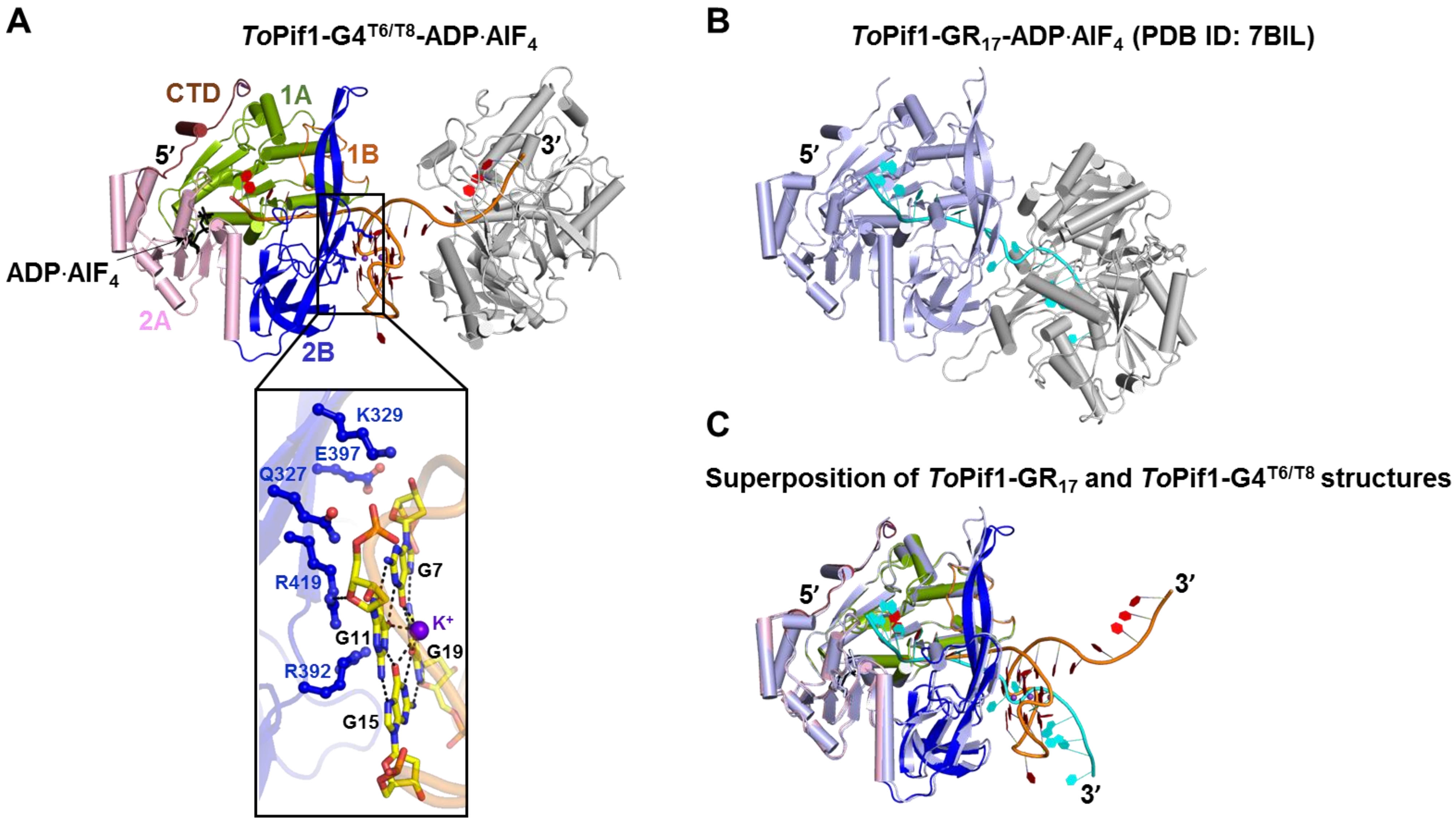


**Supplementary Figure S7. Comparison of domain orientations between the *To*Pif1-G4$^{T6/T8}$-ADP·AlF$_4$ and *To*Pif1-GR$_{17}$-ADP·AlF$_4$ structures. (A)** Cartoon representation of the *To*Pif1-G4$^{T6/T8}$-ADP·AlF$_4$ structure. **(B)** Cartoon representation of the *To*Pif1-GR$_{17}$-ADP·AlF$_4$ structure (PDB: 7BIL (1)). **(C)** Superposition on domain 1A of structures in (A) and (B) with a close-up view of ADP·AlF$_4$.

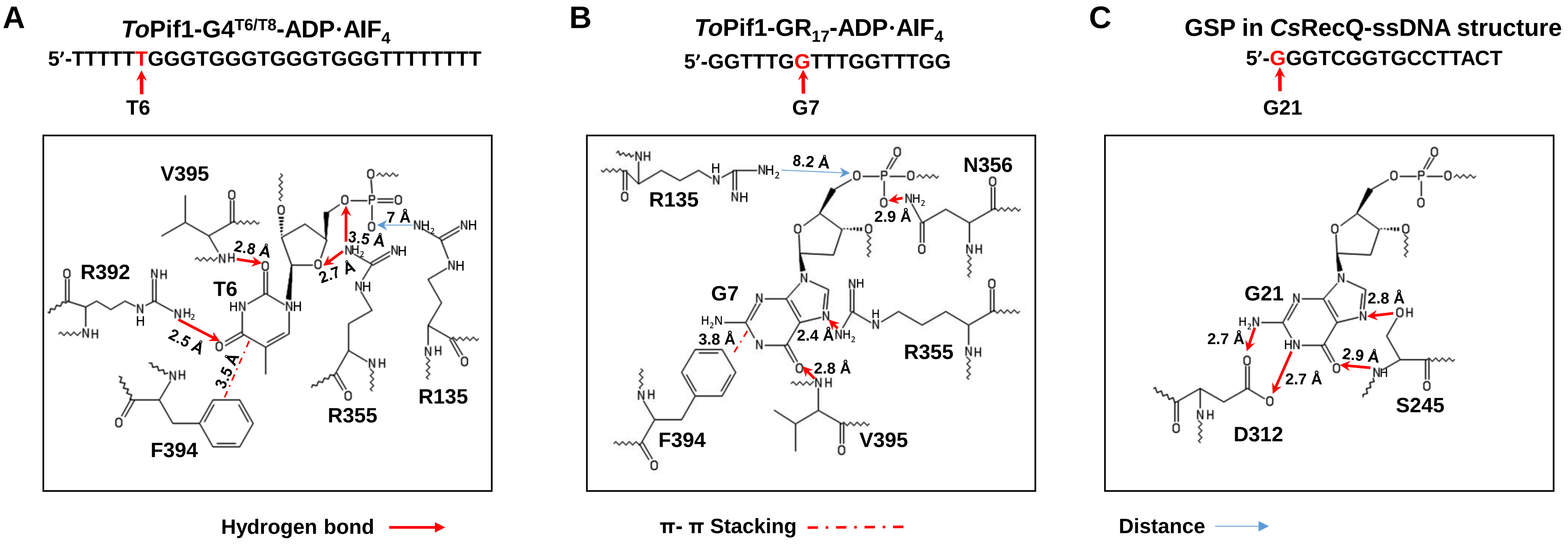


**Supplementary Figure S8. Ligand interaction diagram of T6/G7 in the *To*Pif1 complex and the guanine-specific pocket (GSP) in the *Cs*RecQ-resolved G4 DNA structure.** **(A)** Ligand interaction between T6 and *To*Pif1 in the *To*Pif1-G4$^{T6/T8}$- ADP·AlF$_4$ structure. **(B)** Ligand interaction between G7 and *To*Pif1 in the *To*Pif1-GR$_{17}$- ADP·AlF$_4$ structure (PDB: 7BIL (1)). **(C)** Ligand interaction between G21 and *Cs*RecQ in *Cs*RecQ-ssDNA structure (PDB: 6CRM (4)).

### REFERENCES


1. Dai, Y.X., Chen, W.F., Liu, N.N., Teng, F.Y., Guo, H.L., Hou, X.M., Dou, S.X., Rety, S. and Xi, X.G. (2021) Structural and functional studies of SF1B Pif1 from Thermus oshimai reveal dimerization-induced helicase inhibition. *Nucleic acids research*, **49**, 4129-4143.
2. Zhou, X., Ren, W., Bharath, S.R., Tang, X., He, Y., Chen, C., Liu, Z., Li, D. and Song, H. (2016) Structural and Functional Insights into the Unwinding Mechanism of Bacteroides sp Pif1. *Cell Rep*, **14**, 2030-2039.
3. Lu, K.Y., Chen, W.F., Rety, S., Liu, N.N., Wu, W.Q., Dai, Y.X., Li, D., Ma, H.Y., Dou, S.X. and Xi, X.G. (2018) Insights into the structural and mechanistic basis of multifunctional S. cerevisiae Pif1p helicase. *Nucleic acids research*, **46**, 1486-1500.
4. Voter, A.F., Qiu, Y., Tippana, R., Myong, S. and Keck, J.L. (2018) A guanine-flipping and sequestration mechanism for G-quadruplex unwinding by RecQ helicases. *Nature communications*, **9**, 4201.
